# KatG inactivation generates vulnerabilities in isoniazid resistant strains of *Mycobacterium tuberculosis*

**DOI:** 10.1101/2023.12.07.570702

**Authors:** XinYue Wang, William J Jowsey, Chen-Yi Cheung, Noon E Seeto, Natalie JE Waller, Michael T Chrisp, Amanda L Peterson, Brunda Nijagal, Peter C Fineran, Gregory M Cook, Simon A Jackson, Matthew B McNeil

## Abstract

Drug-resistant strains of *Mycobacterium tuberculosis* are a major global health problem. Resistance to the front-line antibiotic isoniazid is often associated with mutations in the *katG* encoded bifunctional catalase-peroxidase. We hypothesised that perturbed KatG activity would generate collateral vulnerabilities in INH-resistant *katG* mutants, providing new pathways to combat isoniazid resistance. Here, we used whole genome CRISPRi screens, transcriptomics, and metabolomics to generate a genome-wide map of cellular vulnerabilities in a *M. tuberculosis katG* mutant. We discovered that metabolic and transcriptional remodelling compensates for the loss of KatG but in doing so generates vulnerabilities in ribosome biogenesis, and nucleotide and amino acid metabolism. These vulnerabilities were more sensitive to inhibition in an isoniazid-resistant *katG* mutant under *in vitro* and host-relevant conditions and translated to clinical populations. These findings provide an experimental framework for developing novel strategies to combat antimicrobial resistance in *M. tuberculosis* and other bacterial pathogens.

## Introduction

*Mycobacterium tuberculosis* is the primary causative agent of Tuberculosis (TB) and a leading cause of infectious disease morbidity and mortality. Whilst drug-susceptible (DS) strains of *M. tuberculosis* can be treated with a combination of four drugs for six months, drug-resistant (DR) strains require longer treatment regimens with reduced cure rates (1,2). Novel antibiotics and drug combinations that can rapidly sterilise DR-strains and limit the emergence of drug resistance are urgently needed.

The mycobacterial KatG protein is a bifunctional catalase-peroxidase involved in detoxifying reactive oxygen species (ROS) (3–5). Whilst not required for mycobacterial growth *in vitro*, KatG is required in response to ROS and within a murine infection model (6,7). The prodrug isoniazid (INH) is a critical component of front-line anti-tubercular therapy and requires KatG to form the active INH-NAD adduct (8,9). Mutations in KatG that prevent activation of INH are the primary route to INH-resistance (INH^R^), with >70% of INH^R^ clinical isolates having a mutation in *katG* (10). To mitigate fitness costs associated with mutations that enable drug resistance, specific cellular pathways become more important for the growth of DR-strains and consequently more vulnerable to inhibition relative to a drug-susceptible parent (11–15). Prior efforts to identify these collateral vulnerabilities have relied on the systematic screening of antibiotics against DR-strains (16–22). Whilst these studies have identified antibiotics that have increased activity against DR-strains, they have been limited to the use of antibiotics that inhibit only a small number of druggable targets. High-throughput approaches are needed to better define collateral vulnerabilities at a genome-wide level and to identify novel drug targets to prioritise in future drug development programmes.

CRISPR interference (CRISPRi) uses guide RNA (gRNA) sequences and nuclease-deficient Cas9 (dCas9) to transcriptionally silence target genes (23). Whole genome CRISPRi (WG-CRISPRi) offers many advantages as it provides a quantifiable measure of fitness cost or vulnerability associated with target inhibition and allows for the assessment of both non-essential and essential genes (24–28). Here, by combining WG-CRISPRi screens with transcriptional and metabolomic analysis we have generated a genome-wide map of vulnerabilities in an INH-resistant *katG* mutant of *M. tuberculosis*. This experimental framework should accelerate the discovery of new therapeutic approaches for combating DR in *M. tuberculosis* as well as other pathogens.

## Results

### Whole-genome CRISPRi screening to identify genetic vulnerabilities in *M. tuberculosis*

We hypothesised that WG-CRISPRi could be used to identify genes of increased vulnerability in an INH-resistant *katG* mutant of *M. tuberculosis* (KatG^L458QfsX27^, here after referred to as INH^R^-*katG*). To investigate this, we constructed a WG-CRISPRi plasmid library to transcriptionally repress the majority of annotated genes in *M. tuberculosis* (Fig. 1a and Supplementary Table S1). The CRISPRi plasmid pLJR965 encodes a *S. thermophilus* dCas9 mutant (Sth1dCas9) and a gRNA-scaffold sequence, both of which are induced by anhydrotetracycline (ATc) (23,29,30). In total, 22,996 gRNAs were selected to target 3,991 unique protein-coding genes (Supplementary Table S1). Most genes were targeted by 6 individual gRNAs with variation in the predicted PAM strength, gRNA length and predicted gRNA strength (Extended Data Fig.1) (23,25). The WG-CRISPRi library screen was performed in the drug-susceptible (DS)-parent *M. tuberculosis* strain mc^2^6206 by culturing pooled populations for a total of 14 days, with back-dilutions into fresh media with ATc on days 5 and 10 (Fig. 1a). We controlled the level of CRISPRi knockdown by adding different concentrations of the ATc: high, medium, or low, using 300, 30 or 3 ng/ml ATc, respectively. Genomic DNA was harvested on days 5, 10 and 14 and used to generate amplicon libraries for deep sequencing. Amplicon sequencing results demonstrated the number of gRNAs depleted by ≥ 1-log_2_ reduction (i.e. two-fold reduction) in abundance relative to no ATc increased over time with 2,969 gRNAs depleted on at least one of the sampled time points or ATc concentrations (Fig. 1b and Extended Data Fig. 2a). Next, we classified genes as “essential” if ≥ 2 of their targeting gRNAs had ≥ 1-log_2_ reduction in abundance in the day 14 + ATc-300 sampling condition relative to the uninduced (ATc-0) sample. For the DS-parent mc^2^6206, 540 genes meet our essentiality criteria (Fig. 1c). Consistent with prior work showing variations in the time taken to observe a fitness cost when using CRISPRi, the number of genes that had ≥ 2 gRNAs with ≥ 1-log_2_ reduction increased over time (Fig. 1c) (28).

**Figure 1.**
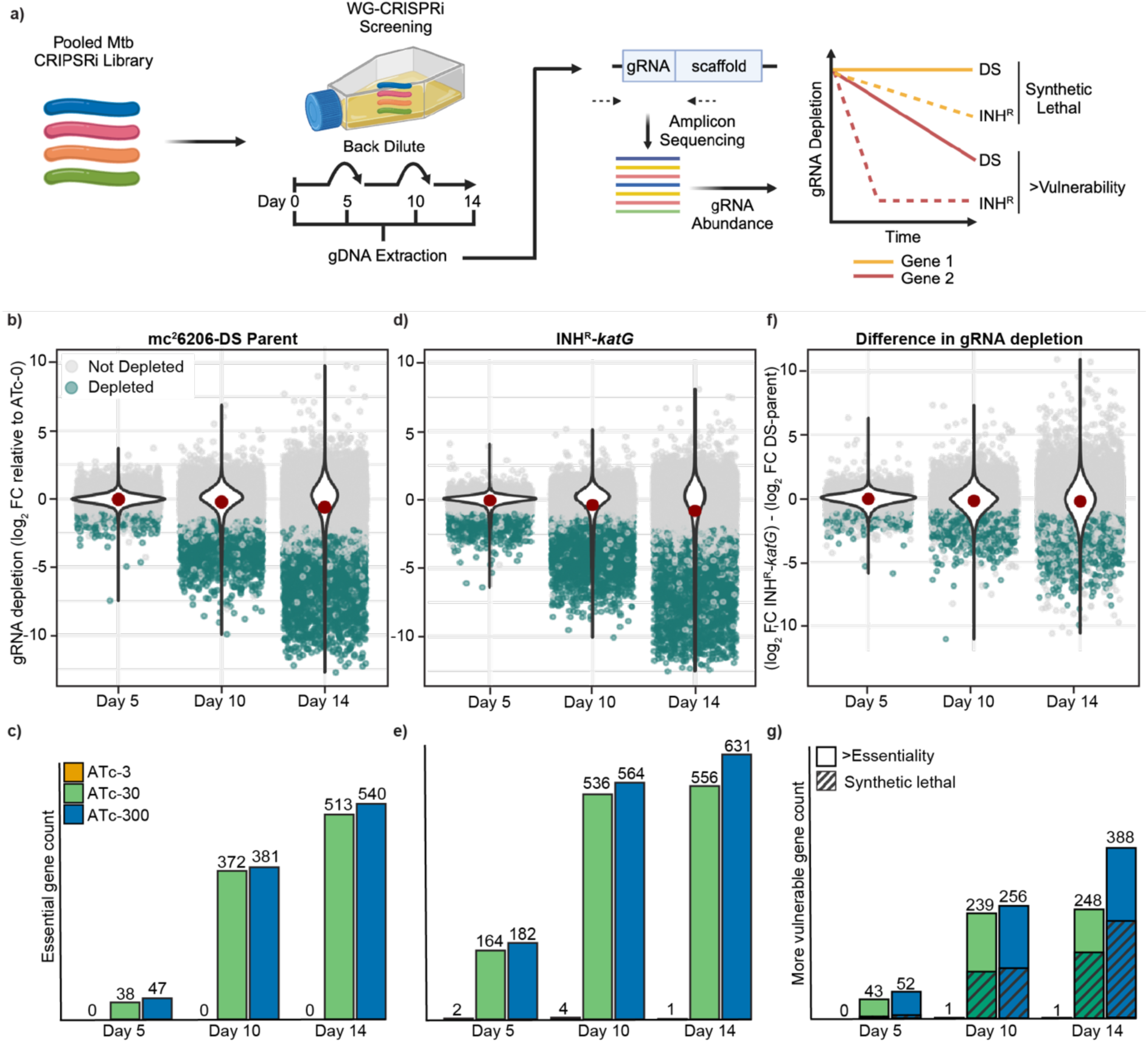
WG-CRISPRi screening in *M. tuberculosis* strain mc^2^6206. **(a)** Experimental design for WG-CRISPRi essentiality screening. The pooled CRISPRi plasmid library is transformed into appropriate *M. tuberculosis* strains. WG-CRISPRi screens were performed for 14 days, with varying concentrations of ATc and at day 5 and 10 back diluted (1/20) into fresh media + ATc to maintain log phase growth. gDNA was extracted, gRNA amplified and sequenced from samples collected on days 5, 10 and 14. The proportion of each gRNA at each ATc concentration within the pooled population was quantified relative to ATc-0 for each sampled time point and plotted on a log2-fold scale as a reduction in gRNA abundance. Increased depletion of gRNA was used to identify genes that are either synthetic lethal or have increased in INH^R^-*katG*. **(b-g)** Summary of gRNA abundance at ATc-300 in the *M. tuberculosis* strain (b) mc^2^6206 DS-parent, (d) INH^R^-*katG*, and (f) the depletion difference between the DS-parent and INH^R^-*katG*. gRNA abundance is relative to the ATc-0 control at each time point. Unchanged gRNAs are coloured grey, whilst gRNA with significant > 2-fold depletion (Benjamini-Hochberg adjusted p-value <0.01) are coloured green. The red dot within each violin plot denotes the mean gRNA depletion. The total number of essential genes identified in the (c) DS-parent, (e) INH^R^-*katG*, and (g) genes identified as being more vulnerable to inhibition in INH^R^-*katG*.

Comparison of our WG-CRISPRi screen to a prior *M. tuberculosis* WG-CRISPRi screen (hereafter referred to as CRISPRi-Bosch) revealed 93.3% agreement in gene essentiality calls, with disagreements for 212 essential and 54 non-essential genes (Extended Data Fig. 2b and Table S2) (25). Of these, most of the 212 disagreed essential genes had previously been defined as low vulnerability in CRISPRi-Bosch (i.e. a vulnerability index (VI) > −5) (Extended Data Fig. 2c-d) (25). Experimental inspection of disagreed genes that we called non-essential but Bosch called essential (i.e. *blaR* and *menE*) demonstrated that they failed to impair the growth of *M. tuberculosis* when grown under conditions comparable to our WG-CRISPRi screen (Extended Data Fig. 2e). The *menE* targeting gRNA did eventually impair growth, but only over a prolonged time course (Extended Data Fig. 2e). Differences in experimental design, including growth media (i.e. OADC vs OADC + Glycerol in CRISPRi-Bosch), the magnitude of gRNA depletion and the timing of essentiality calls (Day 14 vs Day 24 in CRISPRi-Bosch) are likely contributors to the differences in gene essentiality calls. In conclusion, we have developed a WG-CRISPRi platform for *M. tuberculosis* that can reproducibly identify relative differences in the fitness costs associated with targeted gene inhibition.

### WG-CRISPRi identifies genes of increased vulnerability in INH^R^-*katG*

We next used WG-CRISPRi to identify pathways more vulnerable to inhibition in INH^R^-*katG* than the DS-parent strain. We hypothesised that gRNAs targeting pathways of increased vulnerability would manifest as either (i) increased depletion in INH^R^-*katG* at day 14 and/or (ii) be depleted from INH^R^-*katG* earlier in the WG-CRISPRi screen (Fig. 1a). WG-CRISPRi screening identified 3,820 depleted gRNAs and 631 essential genes in INH^R^-*katG* (Fig. 1d-e and Extended Data Fig. 2f). We defined genes as more vulnerable to inhibition in INH^R^-*katG* when (i) they were essential for growth in INH^R^-*katG* and (ii) had ≥2 gRNAs that were depleted by >1-log_2_ fold change than observed in the DS-parent (Fig. 1f -g and Extended Data Fig. 2g). Of the 631 essential genes, we defined 388 genes as “more vulnerable” in INH^R^-*katG* and further classified 168 genes as “synthetic lethal” as they were “essential” in INH^R^-*katG* but “non-essential” in the DS-parent (Fig. 1g).

We hypothesised that gRNAs targeting highly vulnerable essential genes would be maximally depleted at day 14 in the DS-parent and be unable to discern differences in vulnerability. Consistent with this, we identified 38 genes (e.g. *rv0123*, *rplC*, and *rplW*) that meet our criteria of being more vulnerable in INH^R^-*katG* at ≥3 sampling conditions (i.e. earlier time points/lower ATc concentrations) but not at day 14 with ATc-300 (Extended Data Fig. 2h-i). Furthermore, there was no correlation between differential gene expression and altered vulnerability, with only three genes of increased vulnerability being differentially expressed in INH^R^-*katG* (Extended Data Fig. 2j and Table S3). In conclusion, WG-CRISPRi essentiality screens can identify genes that are of increased vulnerability to inhibition in INH^R^-*katG* that cannot otherwise be predicted from changes in gene expression.

### Diverse pathways are more vulnerable to inhibition in INH^R^-*katG*

Having identified more vulnerable genes at a genome-wide level, we sought to identify biological pathways that were enriched for genes of increased vulnerability in INH^R^-*katG*. Pathway enrichment analysis demonstrated that nucleotide metabolism, protein synthesis, respiration and amino acid metabolism were the most enriched functional classes (Fig 2a and Table S4). Although the cell envelope and DNA processing classes were not highly enriched, >20% of genes in the cell wall synthesis and DNA replication subclasses were more vulnerable to inhibition (Extended Data Fig. 3 and Table S4). We hypothesised that gRNAs identified as targeting genes of increased vulnerability in WG-CRISPRi would require less CRISPRi repression (i.e. ATc) to cause a growth impairment in INH^R^-*katG* compared to the DS-parent. To validate the accuracy of our screen we selected 30 gRNAs targeting genes from diverse functional classes for experimental validation. In ATc dose response assays 18 selected gRNAs, including gRNAs targeting *gyrB*, *atpD*, *atpF*, *rpoB*, *hadA, kasA* and *mmpL3,* required less ATc to inhibit the growth of INH^R^-*katG* compared to the DS-parent, (Fig. 2b-d and Extended Data Fig. 4a-d). The gRNAs targeting *clpC1* and *iscS* did not reduce the ATc MIC but altered the shape of the dose response curve (Fig. 2e and Extended Data Fig. 4e). Most gRNAs that required less CRISPRi repression to impair growth also had improved killing against INH^R^-*katG* (Fig. 2f and Extended Data Fig. 4f-m). Although the magnitude of improved killing varied, gRNAs targeting *atpD* and *kasA* were static against the DS-parent yet killed INH^R^-*katG* (Fig. 2f and Extended Data Fig. 4f).

**Figure 2.**
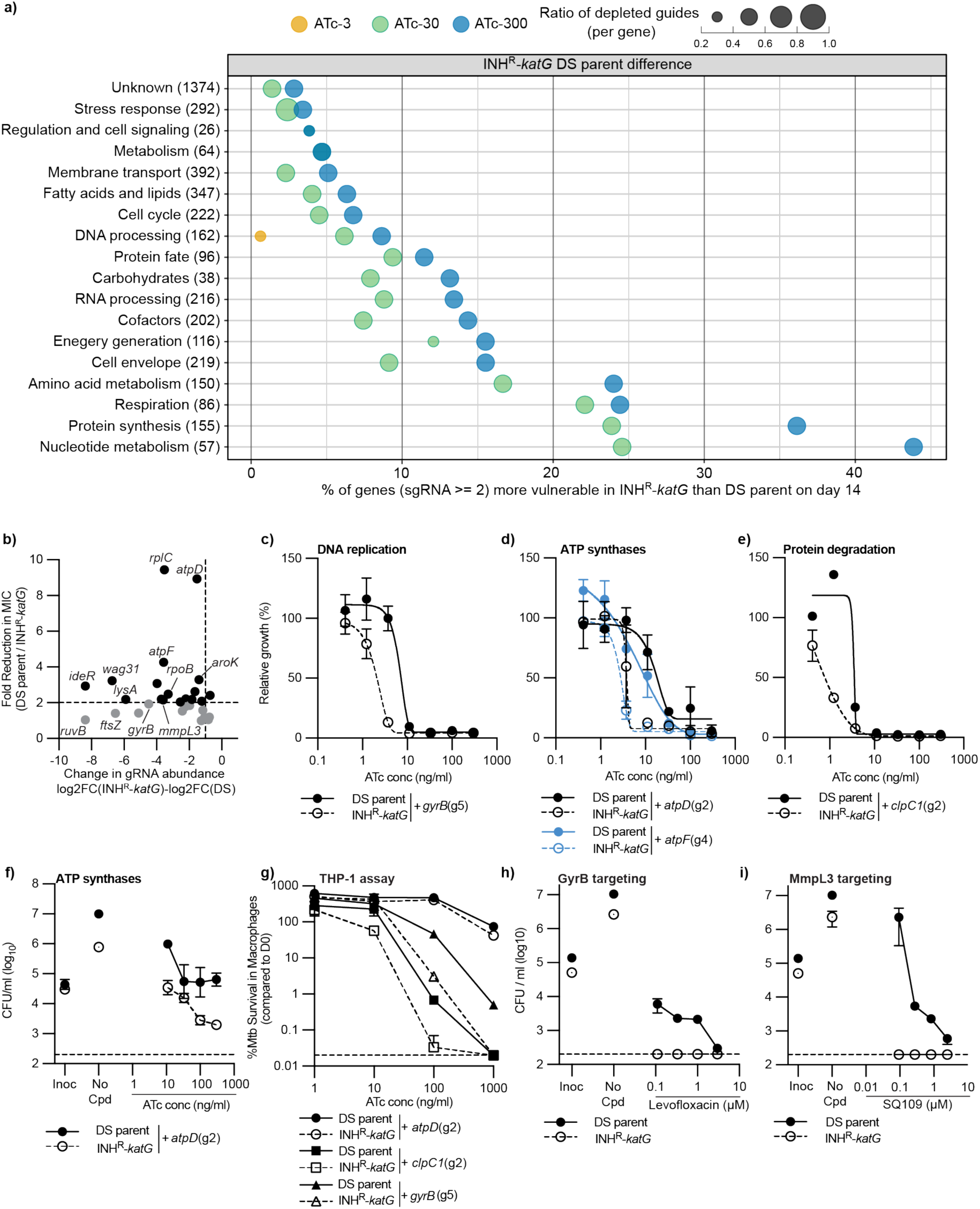
Pathway enrichment analysis defines diverse pathways that are more vulnerable to inhibition in INH^R^-*katG*. **(a)** Pathway enrichment analysis of genes that are of more vulnerable to inhibition in INH^R^-*katG* (i.e. have at least two gRNAs that are significantly depleted) (i.e. Fig 1g). Bubble plot represents data from day 14. Functional classes are described using classifications from the PATRIC database. Within each functional class the size of the bubble indicates the average ratio of gRNAs targeting each gene that is more depleted in INH^R^-*katG.* The colour denotes the ATc concentration from which the amplicon sequencing was performed. The number of genes in each functional class is labelled on the y-axis in brackets. **(b)** Scatter plot for each of the 30 gRNAs that were experimentally validated. Data is plotted as the difference in gRNA abundance between INH^R^-*katG* and the DS-parent from WG-CRISPRi screens (Day14+ATc-300) (x-axis) against the fold change in ATc MIC between INH^R^-*katG* and the DS-parent from ATc dose response assays (y-axis). The vertical dotted line indicates the cut-off for a gRNA to be significantly depleted, whilst the horizontal dotted indicates a two-fold change in ATc MIC. **(c-e)** Growth of *M. tuberculosis* DS-parent and INH^R^-*katG* expressing for gRNA targeting (c) *gyrB*, (d) *atpD* and *atpF* and (e) *clpC1* in ATc dose response assays (mean ± range of two biological replicates, n≥3). The (gx) after each gRNA denotes the specific gRNA targeting each gene. **(f)** Viability plots of *M. tuberculosis* DS-parent and INH^R^-*katG* expressing for gRNA targeting *atpD*. CFU/ml was determined from 96 well plates used in (d). Inoc denotes the starting CFU/ml and no-cpd denotes the detected CFU/ml in the absence of ATc (mean ± range of two biological replicates, n≥3). **(g)** THP-1 macrophage cells were infected with *M. tuberculosis* DS-parent and INH^R^-*katG* cells expressing the stated gRNA. CRISPRi was induced with ATc and intracellular survival was determined as described in the materials in methods (mean ± SD of three biological replicates, n=2). **(h-i)** MBC assays were used to determine the susceptibility of *M. tuberculosis* DS-parent and INH^R^-*katG* to increasing concentrations of (h) levofloxacin and (i) SQ109. Inoc denotes the starting CFU/ml and no-cpd denotes the detected CFU/ml in the absence of compound (mean ± range of two biological replicates, n≥3). Dashed line represents the lower limit of detection.

We hypothesised that if these results translated to a host-relevant model, then gRNAs would have a greater effect on the intracellular survival of INH^R^-*katG* within a THP-1 infection model. Consistent with this, gRNAs targeting *gyrB*, *clpC1* or *rpoB* caused a greater reduction in viable colonies of INH^R^-*katG* compared to the DS-parent (Fig. 2g and Extended Data Fig. 4n) (31). The gRNA targeting *iscS* impaired the intracellular growth of INH^R^-*katG,* whilst the gRNA targeting *atpD* had no effect on the intracellular survival of INH^R^-*katG* (Fig. 2g and Extended Data Fig. 4n). We next hypothesised that if genes of increased vulnerability could serve as collateral drug vulnerabilities, then INH^R^-*katG* would have increased sensitivity to killing by antibiotics that targeted our identified genetic vulnerabilities. Indeed, INH^R^-*katG* was more sensitive to killing by levofloxacin (GyrB targeting), SQ109 and CPD1 (MmpL3 targeting), bortezomib (putative Clp targeting), thiocarlide (HadAB targeting) and BB16F (ATP synthase targeting) (Fig. 2h-i and Extended Data Fig. 5a-d). Except for borezomib, all chemical inhibitors reduced INH^R^-*katG* to the lower limit of detection for all concentrations at or above the DS-parent MIC (Fig 2h-i and Extended Data Fig. 5a-d). Despite the increased vulnerability of *rpoB*, INH^R^-*katG* was not more sensitive to killing by the RNA polymerase inhibitor rifampicin (Extended Data Fig. 5e). In conclusion, INH^R^-*katG* produces collateral vulnerabilities in diverse biological pathways that are more sensitive to inhibition under *in vitro* and host-relevant conditions.

### Alternative redox homeostasis pathways are more vulnerable to inhibition in INH^R^-*katG*

The KatG catalase/peroxidase plays a crucial role in detoxifying hydrogen peroxide (H_2_O_2_) limiting the accumulation of hydroxyl-radicals and subsequent intracellular damage. Consistent with this, INH^R^-*katG* was (i) more sensitive to inhibition and killing by exogenous redox modulators H_2_O_2_, menadione, plumbagin, and ascorbic acid (Fig. 3a-b and Extended Data Fig. 6a-f); and (ii) had a reduced ability to detoxify reactive intermediates following exposure to H_2_O_2_ (Fig. 3c and Extended Data Fig. 6c-d) (32–34). We hypothesised that *M. tuberculosis* would adapt to the loss of KatG catalase/peroxidase by (i) upregulating compensatory detoxification mechanisms or (ii) if compensatory pathways were not upregulated, they would be more vulnerable to inhibition. The majority (∼88%) of genes within the stress response functional subclass were not differentially regulated; with only the universal stress response proteins TB31.7 (*rv2623)*, *rv2624c*, *rv2005c*, *rv2028c*, *HspX (rv2031),* a putative rubredoxin Rv3251c (*rubA*), and the thioredoxin *trxb1* being upregulated, none of which were more vulnerable to CRISPRi (Fig. 3d). The *furA-katG-rv1907c* operon were the most highly upregulated genes, demonstrating INH^R^-*katG* is able to detect increases in oxidative stress (Fig. 3d) (35). Several well-characterized stress-response genes were more vulnerable to inhibition, including *sodA* and *trxB2* yet were not upregulated (Fig. 3d). INH^R^-*katG* was also more sensitive to auranofin a chemical inhibitor of the *trxB2* encoded thioredoxin reductase (Extended Data Fig. 6g-h) (36).

**Figure 3.**
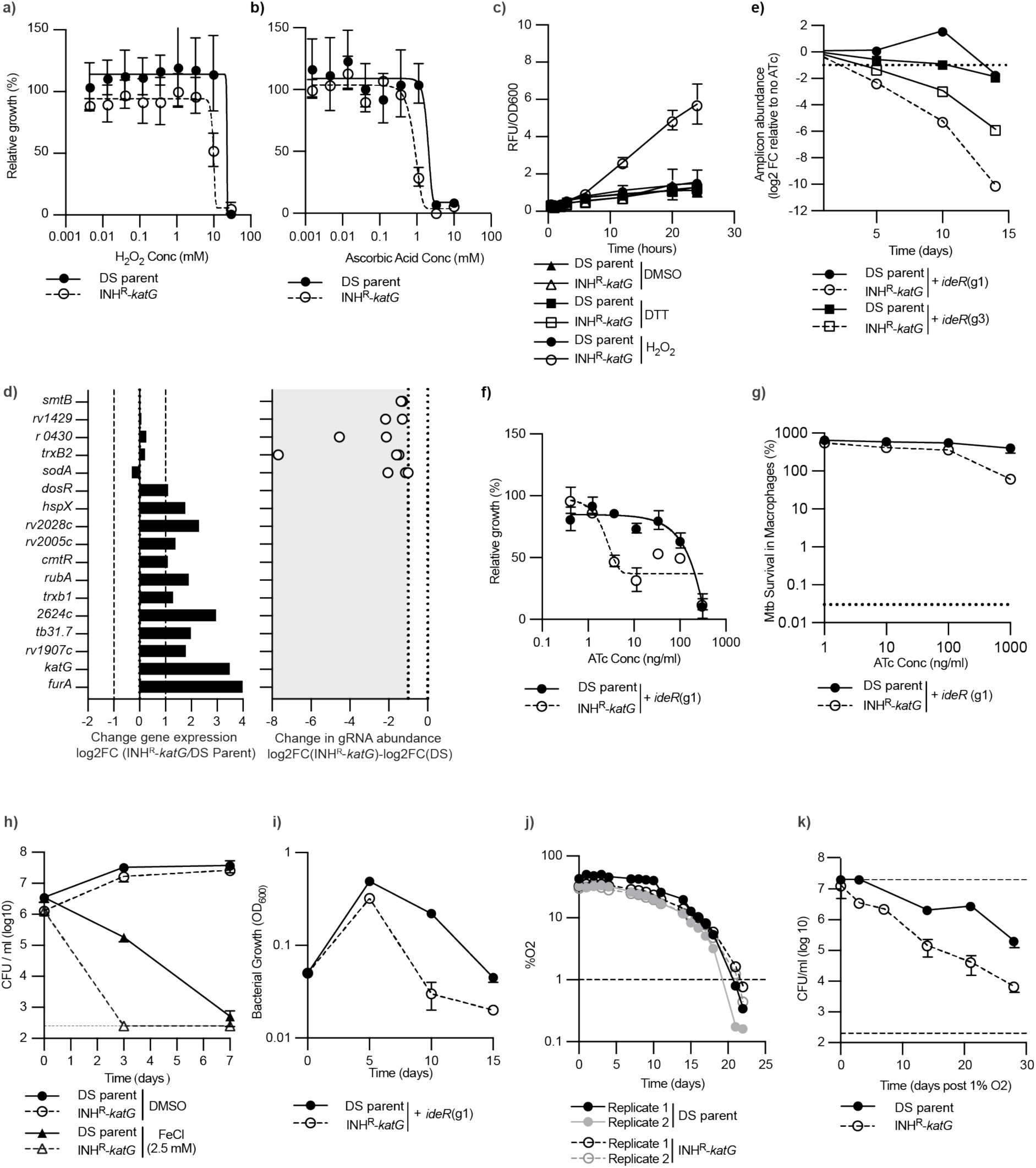
INH^R^-*katG* utilizes alternative redox detoxification pathways to compensate for the loss of *katG*. **(a-b)** Susceptibility of *M. tuberculosis* DS-parent and INH^R^-*katG* to growth inhibition by (a) H_2_O_2_ and (b) ascorbic acid (mean ± range of two biological replicates, n≥3). **(c)** CellRox based detection of reactive oxygen stress in *M. tuberculosis* DS-parent and INH^R^-*katG* when exposed to H_2_O_2_ (mean ± SD of three biological replicates, n=2). **(d)** Changes in gene expression INH^R^-*katG* relative to the DS-parent and increased depletion of gRNA abundance in INH^R^-*katG* relative to DS-parent. Highlighted genes are named using gene name or rv_number. gRNAs that are more depleted in INH^R^-*katG* are presented as white dots, with the positioning of each dot denoting the level of increased depletion on a log2FC scale. Each gene is targeted by multiple gRNA, yet only gRNAs that show a statistically significant depletion on day 14+ ATc-300 are presented. **(e-h)** INH^R^-*katG* is more sensitive to iron dysregulation: (e) Abundance of gRNAs targeting *ideR* throughout the WG-CRISPRi screen as detected by amplicon sequencing. Data is presented for cultures exposed to ATc-300. (f) Growth of *M. tuberculosis* DS-parent and INH^R^-*katG* expressing gRNA targeting *ideR* (mean ± range of two biological replicates, n≥3). The (gx) after each gRNA denotes the specific gRNA targeting *ideR*. (g) Intracellular survival of *M. tuberculosis* DS-parent and INH^R^-*katG* expressing gRNA targeting *ideR* (mean ± SD of three biological replicates, n=2).. (h) Susceptibility of *M. tuberculosis* DS-parent and INH^R^-*katG* to 2.5 mM FeCl as determined by CFU/ml (mean ± SD of three biological replicates, n=2). **(i)** The ability of *M. tuberculosis* DS-parent and INH^R^-*katG* to deplete oxygen during the transition to hypoxia was detected using PreSens oxygen sensing spots. Data represents the individual oxygen consumption curves of two biological replicates from a representative experiment. **(j)** The viability of *M. tuberculosis* DS-parent and INH^R^-*katG* once the concentration of dissolved oxygen was less than 1% was detected by plating for viable colonies.

Interactions between intracellular iron and H_2_O_2_ lead to the formation of hydroxyl radicals (34,37–39). Given the reduced ability of INH^R^-*katG* to detoxify H_2_O_2_ we hypothesised that genes involved in iron storage would be more vulnerable to inhibition. Indeed, our WG-CRISPRi screen identified *ideR*, a transcriptional regulator that regulates intracellular iron storage (25,40,41), as synthetic lethal in INH^R^-*katG* (Fig 3e). Unique gRNAs targeting *ideR* required lower levels of ATc to impair the growth of INH^R^-*katG* in ATc dose response assays, reduced the intracellular survival of INH^R^-*katG* in macrophages and INH^R^-*katG* was more sensitive to killing by exogenous FeCl_3_ compared to the DS-parent (Fig. 3f-h and Extended Data Fig. 6i-j). We further hypothesized that the increased vulnerability of *ideR* would reduce the time needed to observe a fitness cost when using CRISPRi. By back-diluting cultures every 5-days into fresh media + ATc and measuring the maximum growth that was reached, the CRISPRi targeting of *ideR* had an earlier growth inhibitory effect on INH^R^-*katG* compared to the DS-parent. Specifically, growth of INH^R^-*katG* was inhibited by day 10, whilst the DS-parent had only a small fitness cost (Fig. 3i).

In line with a recent transcriptional study of a *M. tuberculosis katG* mutant, we observed an upregulation of DosR and 33 genes (i.e. 65%) in the DosR regulon in INH^R^-*katG* (Table S3) (42). As DosR regulates survival under oxidative stress and hypoxia (43–48), we hypothesised that INH^R^-*katG* would also have an altered ability to survive a hypoxic environment. Despite INH^R^-*katG* consuming oxygen at a comparable rate to the DS-parent and having no reduced viability during adaptation to hypoxia (Fig. 3j), INH^R^-*katG* had a reduced ability to survive under hypoxia compared to the DS-Parent (Fig. 3k). This reduced survivability is consistent with TnSeq experiments and suggests, that similar to mitochondria, hypoxic conditions generate transient bursts in oxidative stress within *M. tuberculosis* (6,49–52). In conclusion, WG-CRISPRi screening reveals that pathways which detoxify or limit the production of DNA damaging hydroxyl radicals become more vulnerable to inhibition in *M. tuberculosis* in the absence of the KatG catalase peroxidase.

### Alterations in amino acid metabolism compensate for perturbed KatG activity

WG-CRISPRi screening revealed that amino acid and nucleotide metabolism were two of the most enriched functional classes for more vulnerable genes in INH^R^-*katG* (Fig. 2a and 4a). We hypothesised that these vulnerabilities were the result of metabolic adaptations to compensate for perturbed KatG activity. Using semi-targeted LC/MS to investigate these adaptations, we identified 127 metabolite peaks, 17 of which were ≥1.5-fold differentially abundant in INH^R^-*katG* (Fig 4b-d and Table S5). Interestingly, the only altered amino acids were those involved in aspartate metabolism (i.e. aspartate, lysine, threonine and methionine), which correlated with genes involved in aspartate metabolism being more vulnerable to inhibition (Fig. 4c-d). Although there was no change in the abundance of aromatic amino acids, genes involved in the chorismate (*aroA, B, G, K, F*) and tryptophan synthesis (*trpA, B, C and E*) were more vulnerable to inhibition in INH^R^-*katG* (Fig. 4d). Aspartate is a precursor to pyrimidine biosynthesis and consistent with this we observed a reduction in N-carbomyl-L-aspartate, the first committed step in pyrimidine biosynthesis, an increased accumulation of downstream pyrimidine (CMP) intermediates and genes involved in early steps of de novo pyrimidine biosynthesis were more vulnerable to inhibition (Fig. 4c). This suggests that altered aspartate metabolism in INH^R^-*katG* in part is needed to maintain an increased demand for nucleotides. The increased accumulation of malate and aspartate, the depletion of oxoglutarate and the increased vulnerability of key steps in the glyoxylate shunt (i.e. *glcB*) suggest that INH^R^-*katG* also reroutes metabolism through the glyoxylate shunt (Fig 4c). These alterations parallel bedaquiline treatment and hypoxia-induced metabolic remodelling in *M. tuberculosis* to bypass the oxidative arm of the TCA cycle, limiting NADH production to reduce electron flow through the electron transport chain and ROS production (53,54).

**Figure 4.**
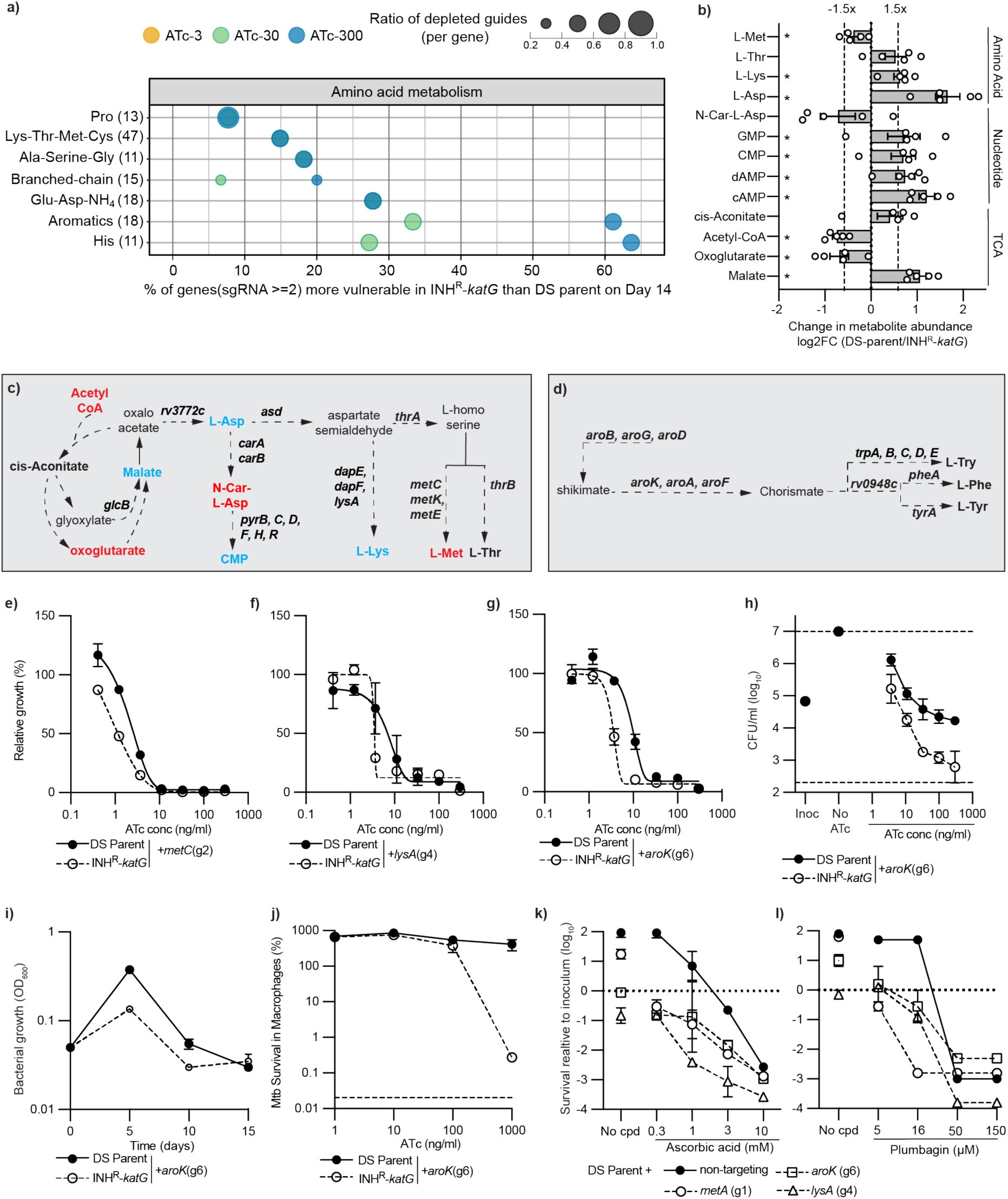
Amino acid metabolism is altered to compensate for the loss of a functional *katG* in *M. tuberculosis*. **(a)** Pathway enrichment analysis of more vulnerable genes as defined by subclasses within the amino-acid class. Subclasses are described using classifications from the PATRIC database. The bubble plot represents data from the day 14. Features of the bubble are as described in figure 2a. Aromatics: Aromatic amino acid metabolism. His: histidine metabolism. **(b)** Changes in the relative abundance of metabolites as determined from five replicates. Metabolites with an asterisk “*” are those that are significantly depleted (i.e. p-value <0.05 as determine by t-test). **(c-d)** Schematics of (c) the metabolic pathways linking aspartate metabolism with the TCA cycle and nucleotide metabolism and (d) aromatic amino acid metabolism. Bolded metabolites highlight those that were detected. Blue, red and black colouring denotes metabolites that are increased, decreased or no-change in INH^R^-*katG*. Bolded genes highlight those that are more vulnerable. **(e-g)** Growth of *M. tuberculosis* DS-parent and INH^R^-*katG* expressing gRNAs that target (e) *metC*, (f) *lysA* and (g) *aroK* in ATc dose response assays (mean ± range of two biological replicates, n≥3). The (gx) denotes the specific gRNA targeting each gene. **(h)** Viability plots of *M. tuberculosis* DS-parent and INH^R^-*katG* expressing for gRNA targeting *aroK.* Inoc denotes the starting CFU/ml and no-cpd denotes the detected CFU/ml in the absence of ATc (mean ± range of two biological replicates, n≥3). Dashed line represents the lower limit of detection. **(i)** Growth kinetics of *M. tuberculosis* DS-parent and INH^R^-*katG* expressing a gRNA targeting *aroK*. Bacterial growth was measured by OD_600_ and back diluted 1/20 into fresh media on days 5 and 10. All strains were grown in 7H9 media with ATc-300 (mean ± SD of three biological replicates, n=2). **(j)** THP-1 macrophage cells were infected with *M. tuberculosis* DS-parent and INH^R^-*katG* cells expressing an *aroK* targeting gRNA. CRISPRi was induced with the stated concentrations of ATc and intracellular survival was determined by plating for viable colonies as described in the materials in methods (mean ± SD of three biological replicates, n=2). **(k-l)** The DS-parent of *M. tuberculosis* mc^2^6206 was pre-depleted for *metA*, *aroK* and *lysA* for 5 days, prior to exposure to (m) ascorbic acid or (n) plumbagin at the state concentrations. No-cpd represents the viability of the pre depleted strains in the absence of compound but in the presence of 300 ng/ml ATc. Data is expressed as the reduction in viable colonies on day 10, relative to the starting inoculum on day 0. A non-targeting gRNA is included as a negative control.

To confirm the increased vulnerability amino acid metabolism, gRNAs targeting methionine (*metA* and *metC*), lysine (*lysA*, and *dapE*), or shikimate synthesis (*aroK* and *aroA*) required less CRISPRi repression (i.e. ATc) to inhibit the growth of INH^R^-*katG* and had an earlier growth inhibitory phenotype against INH^R^-*katG* (Fig 4e-i and Extended Data Fig. 7). The gRNAs targeting *aroK* and *aroG* had improved killing against INH^R^-*katG*, whilst all other gRNAs were static against both strains (Fig. 4h and Extended Data Fig. 8a-i). The gRNAs targeting *aroB* and *aroG* had an earlier inhibitory effect on growth that wasnt observed in ATc dose response assays (Extended Data Fig. 7h-i). Inhibition of *aroK* also impaired the intracellular survival of INH^R^-*katG* in THP-1 infected macrophages (Fig. 4j). Consistent with aspartate and chorismate metabolism compensating for perturbed KatG activity, the depletion of *lysA*, *metA* and *aroK* from the DS-parent increased susceptibility to killing by the redox modulators ascorbic acid and plumbagin (Fig. 4k-l). In conclusion, metabolic remodelling strategies that compensate for perturbed KatG activity in INH^R^-*katG* produce collateral vulnerabilities in amino acid metabolism.

### Ribosome biogenesis is more vulnerable to inhibition in an INH^R^-*katG* mutant

The inhibition of protein synthesis increases oxidative stress and cell death due to disruptions in translational fidelity and increases in misfolded protein levels (55). Consistent with the increased sensitivity of INH^R^-*katG* to oxidative stress WG-CRISPRi screening revealed that >30% of the genes involved in ribosome biogenesis had increased vulnerability in INH^R^-*katG* (Fig. 5a). Confirming the increased vulnerabilities, gRNA sequences targeting *rplC* and *rpsP* required less ATc to inhibit the growth of INH^R^-*katG,* had an earlier growth inhibitory phenotype, and had improved killing against INH^R^-*katG* under *in vitro* conditions and within infected THP-1 macrophages (Fig 5b-e and Extended Data Fig. 9a-b). We hypothesised that if the increased vulnerability of protein synthesis could serve as collateral drug vulnerability, then INH^R^-*katG* would have increased sensitivity to antibiotics that target protein synthesis. Consistent with this, the protein synthesis inhibitors linezolid (oxazolidinone), kanamycin (aminoglycoside) and nitrofurantoin had improved killing against INH^R^-*katG* (Fig. 5f and Extended Data Fig. 9c-d). Linezolid had improved killing against INH^R^-*katG* under *in vitro* conditions, in time-kill assays and within infected THP-1 macrophages (Fig. 5g-h). When used in combination with INH, subinhibitory concentrations of linezolid could also exploit this collateral vulnerability and suppress the emergence of INH resistance (Fig. 5i).

**Figure 5.**
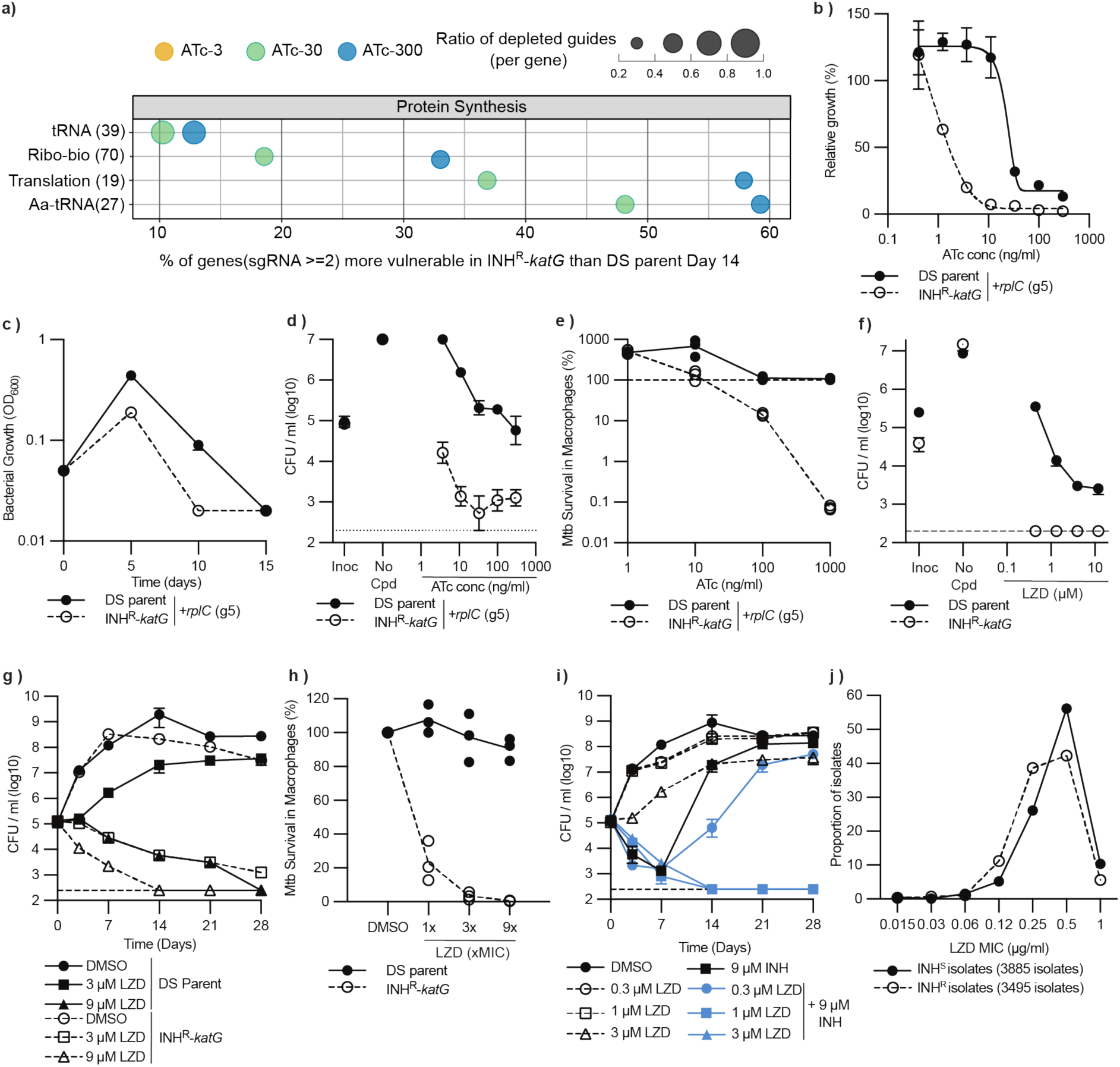
Ribosome biogenesis is more vulnerable to inhibition in an INH^R^-*katG* mutant. **(a)** Pathway enrichment analysis of more vulnerable genes in INH^R^-*katG* as defined by subclasses within the protein synthesis class. Subclasses are described using classifications from the PATRIC database. The bubble plot represents data from the day 14. Features of the bubble are as described in figure 2a. Ribo-bio: Ribosome biogenesis. **(b)** Growth of *M. tuberculosis* DS-parent and INH^R^-*katG* expressing gRNAs targeting *rplC* in ATc dose response assays (mean ± range of two biological replicates, n≥3). The (gx) after each gRNA denotes the specific gRNA targeting each gene. **(c-d)** *M. tuberculosis* DS-parent and INH^R^-*katG* expressing a gRNA targeting *rplC* and assessed (c) in continuous log phase growth in 7H9 media with ATc-300, (d) for viability in ATc dose response assays and **(e)** for intracellular survival within THP-1 macrophage cells (mean ± SD of three biological replicates, n=2). Data presentation is consistent with Figure 4 and experiments were performed as described in material and methods. **(f-h)** Susceptibility of *M. tuberculosis* DS-parent and INH^R^-*katG* to increasing concentrations of linezolid (LZD) as assessed (f) in 96 well plates, (g) in time kill assays and (h) against intracellular M. tuberculosis within THP-1 infected macrophages. For (f) Inoc denotes the starting CFU/ml and no-cpd denotes the detected CFU/ml in the absence of compound (mean ± SD of three biological replicates, n=2). Dashed line represents the lower limit of detection. **(i)** Susceptibility of *M. tuberculosis* DS-parent to LZD or LZD+INH added at the stated concentrations. Dashed line represents the lower limit of detection. **(j)** LZD susceptibility of INH-susceptible (INH^S^) and INH^R^ clinical isolates from the CRyPTIC consortium. Data is expressed as the proportion of isolates that have a specific LZD MIC.

Linezolid was FDA approved for the treatment of MDR-TB as part of a new combination therapy (i.e. BPaL) including bedaquiline and pretomanid. Our data suggests that the excellent efficacy of the BPaL regimen could be attributable to the hyper-susceptibility of INH-resistant (INH^R^) isolates to linezolid, in addition to prior reports of hyper susceptibility to bedaquiline and pretomanid (19,42). To investigate this, we surveyed the CRyPTIC data compendium of ∼12,000 *M. tuberculosis* clinical isolates for altered linezolid susceptibility. Filtering out low quality MIC calls and linezolid resistant phenotypes highlighted that a greater proportion of INH^R^ isolates had a lower linezolid MIC compared to the INH susceptible population (Fig. 5j), with a low yet negative correlation between INH and linezolid susceptibility (Pearson correlation of r = −0.121, p value = 9.89e-26). In conclusion, INH^R^-*katG* is more vulnerable to the inhibition of protein synthesis under *in vitro* and host-relevant conditions, can be exploited by linezolid to suppress INH resistance, and extends into clinical isolates.

## Discussion

Therapeutic strategies that exploit the biological costs of drug resistance have the potential to rapidly sterilize and prevent the emergence of drug resistant pathogens. By applying WG-CRISPRi screens with transcriptional and metabolomic approaches we have built on prior work to generate a genome-wide map of collateral vulnerabilities in an INH resistant *katG* mutant strain of *M. tuberculosis*. This work defines the importance of redox homeostasis in mycobacterial physiology, uncovers how metabolic remodelling compensates for perturbed KatG activity, and describes how newly approved TB combination therapies maybe inadvertently targeting collateral vulnerabilities in DR strains.

This work identified hundreds of genes from diverse biological pathways that are more vulnerable to transcriptional inhibition in an INH resistant *katG* mutant. The enrichment of vulnerable genes to pathways that are known to generate oxidative stress when inhibited is consistent with an increased vulnerability to dysregulated redox homeostasis. Importantly, vulnerabilities identified in our WG-CRISPRi screen were experimentally validated under *in vitro* and host-relevant conditions and could be targeted as collateral drug vulnerabilities. In many instances, the INH^R^-*katG* mutant was more susceptible to killing by gRNAs that targeted more vulnerable genes and was sterilized by chemical inhibitors that otherwise had dose dependent or bacteriostatic killing mechanisms against *M. tuberculosis*. These findings also support prior suggestions that oxidative stress is a critical driver of lethality for many functionally unique antibiotics and highlights the potential of inhibiting redox homeostasis to potentiate antibiotic lethality (56–61).

This work made key functional insights into how changes in the physiology of drug resistant strains generates collateral vulnerabilities. Firstly, disruption of *katG* activity resulted in an increased reliance on alternative ROS detoxification pathways and intracellular iron storage to maintain redox homeostasis. Secondly, metabolic remodelling compensates for perturbed *katG* activity. This included upregulated levels of aspartate to produce increased levels of lysine and maintain methionine production, with each playing a crucial role protecting *M. tuberculosis* against oxidative stress. In *Saccharomyces* species the decarboxylation of lysine into the polyamine cadaverine allows for uncommitted NADPH to be channelled into glutathione metabolism as an antioxidant strategy (62). Consistent with this, increases in cadaverine synthesis have been observed in *M. tuberculosis* INH^R^ mutants using gas chromatography-time of flight mass spectrometry (GS-MS) (63). Whilst we were unable to identify polyamine peaks in our metabolic spectra this may be due to our use of negative mode LC-MS rather than GC-MS (63). Additionally, the increased vulnerability of chorismate metabolism is consistent with recent work showing reduced metabolic flux through this pathway in the absence of KatG (42). Combined, this work reinforces the fundamental role of amino acids in the redox homeostasis (7,64–70). A complementary metabolic strategy is also used to favour the use of the glyoxylate shunt as a strategy to bypass the oxidative arm of the TCA cycle, limit the production of NADH and reduce the flow of electrons through the electron transport chain, thereby limiting ROS byproducts (53,54). Finally, protein synthesis and nucleotide metabolism had the greatest proportion of genes that were more vulnerable to inhibition in INH^R^-*katG*. Both protein synthesis and nucleotide metabolism have known roles in contributing to or disrupting redox homeostasis and have previously been shown to be highly vulnerable genes in *M. tuberculosis* (25). This work extends these findings, highlighting their increased vulnerability in the context of drug resistant strains of *M. tuberculosis* that are experiencing dysregulated cellular physiology and redox homeostasis.

The clinical implications of our findings may be limited by (i) our use of a *katG* frameshift mutant, rather that the KatG^S315T^ mutation that is observed in >70% of INH-resistant clinical isolates and (ii) our use of a *M. tuberculosis* strain that is derived from a single geographic lineage. However, recent reports have extended prior findings of bedaquiline hyper-susceptibility in a *katG* mutant and shown that this (i) translates to a population-level reduction in bedaquiline MIC and (ii) that INH^R^ clinical isolates of *M. tuberculosis* with KatG^S315T^ reproduced the hyper susceptibility of a *katG* deletion mutant against bedaquiline (42). Whilst the population-level reductions against linezolid were small, our results warrant further investigations into how diverse clinical isolates, both susceptible and resistant, respond to killing by linezolid. These combined findings also suggest that the excellent efficacy of the BPaL regimen may be a result of the individual drugs inadvertently targeting collateral vulnerabilities in DR strains of *M. tuberculosis*.

Here we have (i) defined the mechanisms that allow *M. tuberculosis* to adapt to inactivation of *katG* and acquisition of INH-resistance and (ii) defined genes and cellular pathways that as a consequence are more vulnerable to inhibition. Importantly this genetic data translated to existing chemical inhibitors*, ex vivo* models and clinical isolates. In our WG-CRISPRi screen, many of the more vulnerable genes were essential genes and were not detected by transcriptional or metabolic approaches. This highlights the power of WG-CRISPRi in driving novel insights into mycobacterial biology and the identification of highly vulnerable drug targets. The development of portable CRISPRi platforms for diverse pathogens should allow these approaches to be applied to other bacterial pathogens (71–73).

## Materials and Methods

### Bacterial strains and growth conditions

The *M. tuberculosis* strain mc^2^6206 (H37Rv Δ*panCD*, Δ*leuCD*) is an avirulent derivative of H37Rv (74). The isoniazid-resistant *katG* mutant (KatG^L458QfsX27^) used in this study was previously isolated as a spontaneous isoniazid resistant mutant in *M. tuberculosis* strain mc^2^6206 where it was named INH-1 (19). Here, INH-1 is renamed INH^R^-*katG*. *M. tuberculosis* strain mc^2^6206 drug-susceptible parent and INH^R^-*katG* were grown at 37°C in Middlebrook 7H9 liquid medium or on 7H11 solid medium supplemented with OADC (0.005% oleic acid, 0.5% BSA (sigma A-7906), 0.2% dextrose, 0.085% catalase), pantothenic acid (25 µg/ml), leucine (50 µg/ml) and when required kanamycin (20 µg/ml). Liquid cultures were supplemented with 0.05% tyloxapol (Sigma). Throughout the manuscript, 7H9 refers to fully supplemented 7H9 media, whilst 7H9-K refers to 7H9 supplemented media with kanamycin. *Escherichia coli* strain MC1061 was used for the cloning of CRISPRi plasmids. *E. coli* MC1061 was grown at 37°C in LB or on LB-agar supplemented with Functional1.5% agar supplemented with kanamycin at 50 µg/ml.

### Design of *M. tuberculosis* pooled-CRISPRi library

The pooled-CRISPRi library used in this study was designed by sub-setting a published *M. tuberculosis* CRISPRi library that contained 96K gRNAs, targeting 98.2% of *M. tuberculosis* ORFs (25). We selected (i) only gRNAs that target the non-template strand of each ORF and (ii) a maximum of six gRNAs with the strongest PAM scores were selected per gene. In total this CRISPRi library contained 22,996 gRNAs that targeted 3,991 genes. Ten non-targeting gRNA sequences were included as controls. The pooled-CRISPRi library was synthesized and cloned into pJLR965 by Twist Bioscience. Information related to each gRNA in the plasmid pool including gRNA sequence, gene target, abundance in original Twist library, predicted PAM strength, predicted gRNA strength (if available), vulnerability index of target gene based on CRISPRi-Bosch (25), essentiality from prior WG-CRISPRi and TnSeq screens (25,75), product of each gene if known, strand to which the gRNA binds, position of gRNA along the *M. tuberculosis* mc^2^6206 genome and gRNA-ID number in the pool is available in Table S1.

### Transformation and selection of pooled-CRISPRi library into *M. tuberculosis* mc^2^6206

The constructed pooled-CRISPRi library was transformed into the *M. tuberculosis* DS-parent and INH^R^-*katG* as follows. From a glycerol stock, 100 µl of each *M. tuberculosis* strain was used to inoculate 10 ml 7H9 media and grown in a T25 flask until confluent. 100 µl of the outgrowth culture was subcultured into 10 ml 7H9 media and grown until an OD_600_ of 0.4. This was repeated for 12 individual T25 flasks, giving 120 ml of culture for transformation. At an OD_600_ of 0.4, 1 ml of 2 M glycine was added to each T25 and grown overnight. The 120 ml of culture was harvested in three equal volumes, each washed in 10 ml and again in 5 ml of room temperature 10% glycerol. Pellets were resuspended in a combined volume of 10 ml of 10% glycerol. 200 µl of resuspended culture were used to electroporate 5 µl of the constructed pooled-CRISPRi library at a concentration of 50 ng/µl, as previously described (76). Electroporation’s were recovered in 10 ml 7H9 media and grown overnight at 37°C. A minimum of ten transformations were performed to generate a single *M. tuberculosis* CRISPRi library. All recovered transformations were harvested and resuspended to an OD_600_ of 0.4 in 7H9-K media. A 200 µl sample from each transformation was used to determine the efficiency of transformation via serial dilution and plating. From the remaining culture, 2.5 ml was inoculated into 50 ml of 7H9-K in a T-75 flask, giving a starting OD_600_ of approximately 0.02. Cultures were grown at 37°C until they reached an OD of approximately 0.5. The proportion of Kan resistant (transformed) cells in the population was determined at the start and end of the outgrowth and compared to a negative control (untransformed *M. tuberculosis*). At an OD_600_ of 0.5, cultures were harvested, individually resuspended in 5 ml volumes, combined, then the entire library was adjusted to an OD_600_ of 1.0. Cell stocks (1 ml) of the adjusted library were made and frozen at −80°C until required. The total number of Kan-resistant colonies (i.e. transformed) in each library was determined by thawing a single cell stock, performing a tenfold dilution series and plating onto 7H11-K.

### WG-CRISPRi screens

WG-CRISPRi screens were performed in T-25 tissue culture flasks. The screen was initiated by thawing seven 1 ml aliquots of the *M. tuberculosis* pooled-CRISPRi library. The 7 ml of thawed stocks were combined with 7 ml of 7H9-K media, to give an OD_600_ of approximately 0.5. To determine the viability of the thawed library, four separate volumes of 100 µl were removed, diluted and plated for CFUs. To outgrow and expand the *M. tuberculosis* pooled-CRISPRi library, 1 ml of the remaining culture was added to 10 ml 7H9-K media in T-25 tissue flasks and grown for four days to an OD_600_ of approximately 1. Library expansion was repeated in 12 individual T25 flasks. After four days expansion, cultures were combined, harvested and adjusted to an OD_600_ of 1 in 7H9-K. To determine the effects of CRISPRi-mediated gene repression in the pooled-CRISPRi population, 500 µl of OD_600_ 1 adjusted library was added to 10 ml of 7H9-K media in T25 tissue culture flasks with either 0, 3, 30 or 300 ng/ml anhydrotetracycline (ATc) to induce CRISPRi. Five replicate cultures were started for each ATc condition, with this initial inoculation referred to as the day 0 culture. The remaining OD_600_ 1 culture was spun down in two equal volumes, supernatant removed and the cell pellets frozen at –20°C for future gDNA extraction. All T25 flasks were grown at 37°C without shaking for 14 days. To maintain log-phase growth, cultures were back diluted using a 1 in 20 dilution on day 5 and again on day 10 into new 7H9-K media with fresh ATc at the same concentration. Cell pellets with the remaining culture were harvested on day 5, day 10, and day 14 via centrifugation at 4000 rpm for 10 min. The cell pellets were stored at −20°C for gDNA extraction.

### DNA extraction, amplicon library construction and sequencing

Genomic DNA was extracted from the frozen cell pellets using ZymoBIOMICS DNA Miniprep Kits (Zymo Research, #D4300) following the manufacturer’s instructions with the following modifications. Briefly, cell pellets resuspended in 750 µl of ZymoBIOMICs lysis solution were lysed by bead beating for five rounds of 1 min in SPEX SamplePrep MiniG 1600 tissue homogenizer (SPEX, Metuchen, New Jersey, USA) at 1,500 rpm. Samples were eluted with 50 µl of nuclease-free water that was preheated to 60°C. Isolated gDNA was quantified using the Invitrogen Quibt 4 Fluorometer with the Broad Range Qubit kit following the manufacturer’s guidelines (Invitrogen, Carlsbad, CA, USA). The integrity of gDNA was determined by running 5 µl of each gDNA sample on a 1 % agarose gel.

To amplify the gRNA sequence extracted gDNA was diluted to 25 ng/µl and used as a template for PCR amplification. For each sample, 100 ng of gDNA template was used in 50 µl of PCR reaction solution. Master mix was made with Q5 High-Fidelity DNA Polymerase (BioLabs # M0493L) following the manufacturer’s instructions. For each strain 60 individual PCR amplifications were performed (5 replicates of 4 ATc concentration across 3 days). Each PCR reaction contained (i) one of five forward primers that were offset from each other by a single base to dephase the sequencing library and (ii) a reverse primer with no dephasing. We used a dual indexing strategy with ten forward indexes and 12 reverse indexes. Primer sequences are listed in Table S6. PCR conditions were as follows (i) initial denaturation for 2 min at 98°C, (ii) 20 cycles of 98°C for 10 sec, 62°C for 30 sec, and 70°C for 20 sec and (iii) final extension for 2 min at 72°C. PCR products confirmed on a 2% agarose gel, were purified using a GFX PCR DNA and Gel Band Purification Kit (Cytiva, # 28903471) and quantified using the Invitrogen Qubit 4 Fluorometer with the High Sensitivity Qubit kit following the manufacturer’s guidelines (Invitrogen, Carlsbad, CA, USA). Each of the 60 purified PCR products per strain were normalized, pooled, then size-selected (245–255 bp) using a Pippin Prep (Sage Science) with 2% agarose dye-free gel cassette (#CEF2010, Sage science, Beverly, USA). The resulting libraries were quantified by Qubit 4 fluorometer (Invitrogen, Carlsbad, CA, USA), then sequenced (100 bp single-end; Illumina NextSeq 2000 at SeqCenter (https://www.seqcenter.com/).

### Deep sequencing data analyses and hit calling

Demultiplexed fastq files were converted to tabular format using SeqKit (77). Sequence features (promoter, gRNA and scaffold sequences) were identified using stepwise fuzzy string matching regardless of the sequencing quality scores (since the expected promoter and gRNA sequences are known, low-quality base calls do not negatively impact downstream analyses). First, only reads with identified promoter regions (upstream of the expected gRNA sequence) were included in subsequent analyses (typically > 99% of raw reads). To account for variable gRNA sequence lengths in the CRISPRi pool, we then used the 34 nt starting from the +1 site as a unique sequence ‘tag’ to match with the expected gRNA sequence pool (Table S1). Only tags with perfect matches to the expected gRNA pool were progressed to generate gRNA count tables for subsequent analyses. Differential abundance tests were performed via the edgeR (v3.42.4) package (classic exact test) to analyse the foldchange of gRNA abundance at each timepoint relative to the corresponding ATc-0 samples (78).

Essential genes were defined when at least 2 gRNAs targeting the same gene of interest had at least a two-fold change in gRNA abundance (Benjamini-Hochberg adjusted p-value < 0.01) relative to the ATc-0 control day 14 with ATc-300. Genes were identified as being more vulnerable to inhibition in INH^R^-*katG* when the gene of interest was (i) classified as essential in INH^R^-*katG* and (ii) had ≥2 gRNAs that were depleted in INH^R^-*katG* by >1-log_2_ fold relative to the depletion of the same gRNA in the DS-parent at day 14 with ATc-300. All data processing scripts are available from GitHub (https://github.com/Cecilia-Wang/2023_CRISPRI)

Functional classification of each target gene was conducted using the PATRIC database from the bacterial and viral bioinformatics resource center (79). Class and subclass information of each gene was obtained via the subsystems table of the H37rv PATRIC database. In cases where functional classification was missing for a gene, manual classification was applied based on its end products and biological processes.

### Construction and transformation of CRISPRi plasmids that target genes of interest

For validation studies, gRNA sequences of interest from the pooled-CRISPRi library were cloned as individual CRISPRi plasmids, as previously described (29,30,76,80). Briefly, the gRNA sequence and a complementary sequence were ordered with GGGA and AAAC overhangs. Oligos were annealed and cloned into the CRISRPi plasmid pLJR965 using BsmB1 and confirmed using sanger sequencing as previously described (76). All ordered oligos and constructed CRISPRi plasmids are listed in Table S7. CRISPRi plasmids were transformed into *M. tuberculosis* DS-parent or INH^R^-*katG* as previously described (76).

### CRISPRi phenotypic assessment of essentiality and viability in 96 well plates

To determine the consequences of targeted gene repression on bacterial growth and viability, phenotypic assays were performed as previously described (30,76). Briefly, ATc dose response assays were performed in 96 well plates. *M. tuberculosis* mc^2^6206 strains containing CRISPRi plasmids grown in 7H9-K and diluted to an OD_600_ of 0.01 in 7H9-K in a deep well 96 well plate. 96 well assay plates were prepared with a 3-fold dilution of ATc along the Y-axis starting at 300 ng/ml of ATc in row H with a starting inoculum of OD_600_ 0.005. This was achieved by adding 75 µl of 7H9-K to all wells of columns 3–10 except row H. 113 µl of 7H9-K containing the starting concentration of ATc (i.e., 600 ng/ml ATc) was added to row H of columns 3–10. ATc was diluted along the vertical axis, transferring 37.5 µl between columns, up to row B. Row A was used as a no ATc control. Columns 1, 2, 11 and 12 contained 150 µl of 7H9-K as contamination and background controls. Seventy five µl of OD_600_ adjusted culture was added to each well to achieve a starting OD_600_ of 0.005. Each column represents the ATc dilution gradient for a single *M. tuberculosis* strain containing a unique CRISPRi plasmid. All experiments included a nontargeting sgRNA (i.e., pLJR965) as a negative control. To assess the fitness costs of gRNAs on growth, duplicate plates were grown at 37°C without shaking for 10 days. OD_600_ was measured using a Varioskan-LUX microplate reader. OD_600_ reads from duplicate plates relative to the growth of the no-ATc control were analysed using a nonlinear fitting of data to the Gompertz equation (29).

To assess the effects of gRNAs that targeted more vulnerable genes on bacterial viability, duplicate 96 well assay plates were set up as described above. Viability at day 0 was determined using a 4-point ten-fold dilution of the 0.01 diluted culture, with 5 µl of each dilution spotted onto to 7H11-K agar plates. At Day 5, culture from rows A and D-H were transferred to a new 96 well plate to be diluted. A 4-point ten-fold dilution gradient was performed and 5 µl of each dilution was spotted onto to 7H11-K agar plates. Plates were incubated at 37°C for 4–5 weeks and colonies were counted.

### CRISPRi phenotypic assessment of essentiality under continuous log phase growth

The increased vulnerability of target genes were also assessed using growth curves in which culture was back-diluted to maintain a continuous log phase growth. Experiments were performed by diluting *M. tuberculosis* mc^2^6206 strains containing CRISPRi plasmids in 7H9-K to an OD_600_ of 0.5. 1 ml of culture was added to 10 ml of 7H9-K in a T25-flasks with ATc-300 to a starting OD_600_ of 0.05. Cultures were grown without shaking at 37°C. At day 5, the OD_600_ of culture was determined and 0.5 ml of culture was back-diluted into 9.5 ml of 7H9-K in a T25-flasks with ATc-300 and grown without shaking at 37°C. This was repeated on day 10, with the final OD_600_ being determined on day 15.

### Compound susceptibility and viability assays

The susceptibility of the DS-parent or INH^R^-*katG* to different compounds was determined using Minimum inhibitory concentration (MIC) assays as previously described (19,81). Briefly, inner wells (rows B–G, columns 3–11) of a 96-well flat-bottomed microtiter plate (ThermoFisher Scientific) were filled with 75 µl 7H9 media. Outer wells were filled with 150 µl 7H9 media and left as media only controls. 113 µl of 7H9 media containing compound of interest at the required starting concentration was added to column 2 of row B-G. Compound was diluted 3-fold, by transferring 37.5 µl between wells, down to column 10. Column 11 was kept as solvent only. Strains were diluted to an OD_600_ of 0.01. Seventy-five µl of diluted culture was added to inner wells of the 96-well flat-bottomed microtiter plate containing compound to achieve a starting OD_600_ of 0.005 in a final volume of 150 µl. Plates were incubated at 37°C for 10 days without shaking. After 10 days, plates were covered with plate seals, shaken for 1 min and the OD_600_ was determined using a Varioskan Flash microplate reader (ThermoFisher Scientific). OD_600_ reads from duplicate plates were corrected for background, and values relative to the growth of the no-ATc control were analysed using a nonlinear fitting of data to the Gompertz equation. Assays to determine bacterial viability in response to compound exposure were set up as described above, with viability determined on days on days 0 and 10. Viability at day 0 was determined using a 4-point ten-fold dilution of the 0.01 diluted culture, with 5 µl of each dilution spotted onto to 7H11 agar plates. At Day 10, culture was removed from appropriate wells and transferred to a new 96 well plate to be diluted. A 4-point ten-fold dilution gradient was performed and 5 µl of each dilution was spotted onto to 7H11 agar plates. Plates were incubated at 37°C for 4–5 weeks and colonies were counted.

### Time kill experiments

Time kill experiments were performed using previously established protocols (19,82). Briefly, cultures were diluted to an OD_600_ of 0.1 in 7H9 media, with 500 µl added to 9.5 ml 7H9-supplemented media in a T25 flask. 50 µl of diluted compounds were added, with DMSO at a final concentration of 0.5%. In co-treatment experiments, antibiotics were added so the final concentration of DMSO was ≤1%. Culture was removed on stated days, diluted, and spotted as described above for MBC assays to determine the number of viable colonies.

### Compound susceptibility and viability assays against *M. tuberculosis* strains pre-depleted for genetic targets

*M. tuberculosis* strains depleted for genes of interest using CRISRPi were prepared by diluting log phase culture to an OD_600_ of 0.005 in 10ml 7H9-K with 300 ng/ml ATc. Cultures were grown without shaking for 5-days at 37°C to pre-deplete target genes. After 5-days *M. tuberculosis* expressing a non-targeting control gRNA was diluted 1/10 into 2 ml of 7H9-K+ATc in a deep well 96 well plate to a theoretical OD_600_ of 0.01. As the transcriptional inhibition of *metA*, *lysA* and *aroK* in this study inhibits bacterial growth, 2 ml of undiluted culture was added directly to a deep well 96 well plate at theoretical OD_600_ of 0.01. Assay plates for susceptibility assays were prepared as described above in 7H9-K+ATc. Seventy-five µl of culture from the deep well plate was added to assay plates as described above. Viable colonies were determined on day 0 and 10 as described above. Plates were incubated at 37°C for 4–5 weeks and colonies were counted. Data is present as the change in CFU/ml relative to the inoculum.

### Hypoxia survival experiments

Oxygen sensing spots (PreSens, Germany) were adhered to the inside of 100 ml glass vials and sterilized before use (83). Vials were inoculated with 1 ml *M. tuberculosis* adjusted to an OD_600_ of 1 into 29 ml 7H9 for a starting concentration of OD_600_ ∼0.03. Glass vials were stopped with a rubber stopper to prevent gaseous exchange. Cultures were incubated at 37°C with shaking (200 rpm). The oxygen concentration was measured following the manufacturers guidelines by reading the sensor spot through the outside of the flask using a fibre optic cable connected to a Fibox 4 oxygen meter (PreSens). CFU samples were taken using a hypodermic needle that was inserted through the rubber stopper to remove 500 µl culture. Samples were diluted along a four point ten-fold dilution series, spotted onto 7H11-K and incubated at 37°C. Colonies were counted once visible growth was detected, i.e. approximately 4 weeks

### CellROX measurements of oxidative stress

CellROX measurements were performed following published protocols (84). Briefly, mid-log phase cultures of *M. tuberculosis* mc^2^6206 DS-parent or INH^R^-*katG* were harvested by centrifugation (4000 rpm, 10 minutes), washed, resuspended in sterile phosphate buffered saline and diluted to an OD_600_ of 1.0. 100 μL of diluted culture was added to black, clear bottom 96-well microtiter plates (Thermofisher #165305). Culture was then treated with 1 mM of hydrogen peroxide (H_2_O_2_), dithiothreitol or the solvent control DMSO. Plates were incubated at 37°C for 1 h. CellROX Green reagent (Thermofisher #C10444) was added to the desired wells at a final concentration of 5 μM and was returned to the incubator in the dark for 30 min. OD_600_ and fluorescence (λEx 485 nm/λEm 520 nm) was measured using using a Varioskan Flash microplate reader (ThermoFisher Scientific).

### Macrophage infection assays

THP-1 macrophage infection studies were performed using previously described protocols (31). Briefly, the human monocytic cell line THP-1 (ATCC Cat# TIB-202) was cultured in standard RPMI 1640 macrophage medium supplemented with 10% inactivated fetal bovine serum and 1 mM sodium pyruvate at 37°C with 5% CO_2_. THP-1 monocytes (5 × 10^5^ cells/well) were differentiated overnight using 100 ng/ml phorbol myristate acetate (PMA) and seeded in a 24 well-plate. Differentiated macrophages were infected with a mid-logarithmic phase culture of *M. tuberculosis* with or without a CRISPRi plasmid (OD 0.4–0.8) at a multiplicity of infection (MOI) of 10:1 (10 bacteria/1 cell). Infection was allowed to proceed for 1 h. Cells were then washed 3 times with pre-warmed complete RPMI to remove extracellular bacilli. RPMI media containing supplements (pantothenic acid 25 µg/ml and leucine 50 µg/ml), 0.1% BSA and either antibiotic or ATc at varying concentrations were added to the infected cells and incubated at 37°C with 5% CO_2_. After 3 days, infected cells were lysed in distilled water containing 0.1% tyloxapol for 5 min at room temperature to determine the number of CFU/ml on 7H11 agar. For strains with CRISPRi plasmids 7H11 was supplemented with Kan. The percentage of cell viability was determined by normalizing CFU/ml counts at day 3 following compound treatment relative to inoculum as determined on day 0.

### Metabolite extraction, mass spectrometry, and semi-targeted analyses

Metabolites were extracted from cultures of both DS-parent and INH^R^-*katG* as follows. From glycerol stocks, 150 µl of each strain was used to inoculate 10 ml of 7H9 and were grown in T25 flasks at 37°C without shaking until confluent. One hundred µl of these cultures were then used to inoculate T25 flasks containing 10 ml of fresh 7H9 medium. Once confluent (approximately 2 weeks), these cultures were used to inoculate 6 ml of fresh 7H9 to a density of OD_600_ 0.25 and grown for a further ∼48 h (OD_600_ of 0.5-1.2) in T25 flasks at 37°C without shaking. Culture volumes equivalent to 5 ml of culture at an OD_600_ of 1 were filtered through 0.22 µm filters (Millipore, # GVWP02500) via vacuum filtration. Cell-laden filters were suspended in 2-ml bead beater tubes (SSIbio, # 21276) containing 1 ml of fresh metabolite extraction solvent (2:2:1 ratio of acetonitrile (Sigma-Aldrich, # 900667), methanol (>99.8%), distilled and deionised water (18.2 Ω)) and ∼200 µl of 0.1 mm silica beads (dnature, # 11079101z), and cells were lysed by bead beating at 4000 rpm three times for 30 s. Samples were rested for 30 s on dry ice (solid CO_2_) between each bead beating run. Cell lysates were centrifuged at 13000 ×g for 10 min at 4°C, then ∼400 µl of the soluble fraction were transferred to 0.2 µm Spin-X filter columns (Costar, # 8196) and centrifuged at 13000 ×g for 3 min at 4°C. Filtered lysates were transferred to fresh, pre-chilled microcentrifuge tubes before storage at −80°C. Replicates of the DS-parent and INH^R^-*katG* were prepared in parallel. Five replicates were prepared in total, with each replicate prepared on separate days. Samples were shipped on dry-ice to Metabolomics Australia (University of Melbourne, Victoria, Australia) for Mass Spectrometry (MS) analysis of the metabolite. Samples were run alongside an in-house standard library containing 550 polar metabolites that were used as references for the assignment of DS-parent and INH^R^-*katG* sample metabolite peaks, resulting in a semi-targeted approach. Metabolites were separated and detected using a Vanquish Horizon UHPLC system (Thermo Scientific) coupled to an Orbitrap ID-X Tribrid mass spectrometer (Thermo Scientific). Chromatography conditions were performed as previously reported (85) with modifications. Briefly, separation was performed using a SeQuant *zic*-pHILIC column (150 mm × 4.6 mm, 5 µm particle size; Merck) at 25°C, with a binary gradient of solvent A (20 mM ammonium carbonate (pH 9.0; Sigma-Aldrich) and solvent B (100% acetonitrile (Merck, # 100029). The gradient of A/B solvents was run at a flow rate of 300 μl/min as follows: 0.0 min, 80% B; 0.5 min, 80% B; 15.5 min, 50% B; 17.5 min, 30% B; 18.5 min, 5%; 21 min, 5% B; 23–33 min, 80% A. For metabolite detection, the Orbitrap ID-X Tribrid Mass Spectrometer was coupled to a heated electrospray ionisation source and performed as follows: sheath gas flow 40 arbitrary units, auxiliary gas flow 10 arbitrary units, sweep gas flow 1 arbitrary units, ion transfer tube temperature 275°C, and vaporizer temperature 320°C. The radio frequency lens value was 35%. Data was acquired in negative polarity with spray voltages of 3500 V. Samples were run in a random order, and the quality of data produced was assessed by the peak variation of pooled samples (all samples combined equally) and four internal standards (^13^C_5_,^15^N_1_ Valine, ^13^C_6_ Sorbitol, ^13^C,^15^N-UMP, ^13^C,^15^N-AMP) that were added to each sample. The data was collected using Thermo Tracefinder (V 4.1) (General Quan Browser). Metabolites were assigned to sample peaks in El-Maven v.0.12.1 by comparison to the peaks in the standard library. WT and INH^R^-*katG* peaks that were also observed in metabolite extraction solvent-only samples (at >20% of the WT) were excluded. Identified metabolites of the DS and INH^R^-*katG* samples were provided as raw values from the area under the curve. Raw data were processed and analysed using the MetaboAnalyst v5.0 web server (https://www.metaboanalyst.ca/docs/About.xhtml) as follows (86). Peak intensities of all identified metabolites in a combined dataset of DS-parent and INH^R^-*katG* were processed using the Statistical Analysis [one factor] pipeline. Here, the dataset was median normalised and log_10_ transformed. Metabolites with fold changes >1.5× (p<0.1) between the DS and INH^R^-*katG* groups were considered to be altered in relative abundance.

### RNA extraction and analysis of RNA sequencing

Five replicate cultures (of both DS-parent and INH^R^-*katG*) were inoculated into 7H9 media at a starting OD_600_ of 0.1 in T25 tissue culture flasks, grown for 3 days at 37°C without shaking, then harvested and RNA extracted as previously described (76). Briefly, the volume of culture harvested was determined as follows (OD_600_ x volume of culture(ml) = 2.5). Harvested pellets were resuspended in 1 ml TRIzol, bead beaten in a 2 ml tube with 200 µl of 0.1 mm Zirconia/silicon beads for 3 cycles of 30 sec at 4800 rpm followed by 30 secs on ice and frozen overnight at −20°C. Frozen samples were thawed, mixed with 0.2 ml chloroform and centrifuged in Invitrogen PhaseMaker tubes (Cat No: A33248) for 15 minutes at 12,000 g. The clear, upper aqueous phase was transferred to a clean 1.5 ml Eppendorf tube and mixed with an equal volume of ethanol. RNA was extracted using Zymo-RNA Clean and Concentrator (Cat No: R1019) and DNase treated using Invitrogen Turbo DNA-free kit (Cat No: AM1907). Removal of DNA was confirmed by PCR using 1 µl of extracted RNA as a template with the primer combination of MMO200+MMO201. Extracted RNA was quantified and quality controlled using the Aligent 2100 Bioanalysis system following the manufacturer guidelines. RNA was prepared using GenTegra RNA tubes (#GTR5025-S) and shipped at room temperature to SeqCenter for RNA sequencing (PRJNA1041353).

Each replicate was sequenced using the 12M Paired End rRNA depletion RNA sequencing service. Library preparation was performed with Illumina’s Stranded Total RNA Prep Ligation with Ribo-Zero Plus kit and 10bp IDT for Illumina indices. Sequencing was done on a NextSeq2000 giving 2 x 51bp reads. Demultiplexing, quality control, and adapter trimming was performed with bcl-convert (v3.9.3). Adaptor removal and quality trimming were conducted with bbduk (a part of the BBTools suite https://sourceforge.net/projects/bbmap/). In all cases, low-quality reads with a Phred quality score <10 were fremoved. Contaminant removal was then carried out to filter out all reads that have a 31-mer match to PhiX (a common Illumina spikein) allowing one mismatch. After pre-processing, 13,719,397 read pairs on average per sample were used for downstream analyses.

The cleaned paired-end transcriptomic FASTQ files were then aligned to the *Mycobacterium tuberculosis* mc^2^6206 complete genome (NCBI Accession number: PRJNA914416) with Bowtie2 using the default settings (v2.4.5) (87). The output alignments were saved as SAM files, converted to sorted BAM files, and produced index BAI files with SAMtools (v1.16.1) (88). The resulting alignment files (i.e. BAM and BAI files) were loaded in R (v4.3.0) with the package Rsamtools (v2.16.0) (89,90). Gene counts were calculated using packages GenomicFeatures (v1.52.0) and GenomicAlignments (v1.36.0). Differential expression of each gene was calculated with DESeq2 (v1.40.1) between DS-parent and INH^R^-*katG* strains. Gene expression with at least 2-fold difference (Benjamini-Hochberg adjusted p-value <0.05) between DS and INH^R^-*katG* were considered to be differentially expressed. Scripts of all data processing are accessible from GitHub.

### Analysis of CRyPTIC Consortium data

The CRyPTIC consortium list of 12,288 strains and associated MIC against 13 antibiotics (CRyPTIC_reuse_table_20221019.csv) was obtained from the CRyPTIC consortium (91) (http://ftp.ebi.ac.uk/pub/databases/cryptic/release_june2022/reuse/). Data were filtered to remove strains with either a “LOW” or “NA” quality INH or linezolid phenotype. MIC data that were listed as being greater than a specific value (e.g. “>2”) were rounded up to the next value on a two-fold dilution curve (i.e. MIC of >2 becomes 4). MIC data specified as “<=” were treated as having an MIC of the specified value (i.e. MIC of <=0.025 becomes 0.025). INH-resistant (INH^R^) strains were designated as those having a CRyPTIC consortium resistant (R) phenotype. INH^R^ were further filtered into high level INH-resistant strains with an MIC >= 0.8 µg/ml, to remove potentially low level non *katG* mediated resistance mechanisms. INH-susceptible (INH^S^) strains were taken as having a CRyPTIC consortium susceptible (S) phenotype. As linezolid is only approved for the treatment of MDR strains of *M. tuberculosis* and to avoid this confounding the identification of differences in linezolid susceptibility we removed linezolid resistant strains (MIC >= 2 mg/ml) from both populations. Correlation between the INH and LZD MIC of the combined INH^R^ and INH^S^ population dataset was determined using a Pearson Correlation.

## Acknowledgments

This research was supported by the Health Research Council of New Zealand via project grant 20/459 and Sir Charles Hercus Health Research Fellowships to MM and SAJ (grants 22/156 and 23/228, respectively). NJW was supported by a University of Otago Doctoral Scholarship. We thank members of the Department of Microbiology and Immunology 6^th^ floor for helpful discussions.

## Author Contributions

MBM and SAJ conceived the project. XW, NJW, WJ, CYC, MC, NS, SAJ and MBM performed the experiments. XW, WJ, GMC, PCF, SAJ and MBM interpreted the data. XW and MBM wrote the manuscript with input from all other authors.

## Extended Data

**Extended Data Fig. 1.**
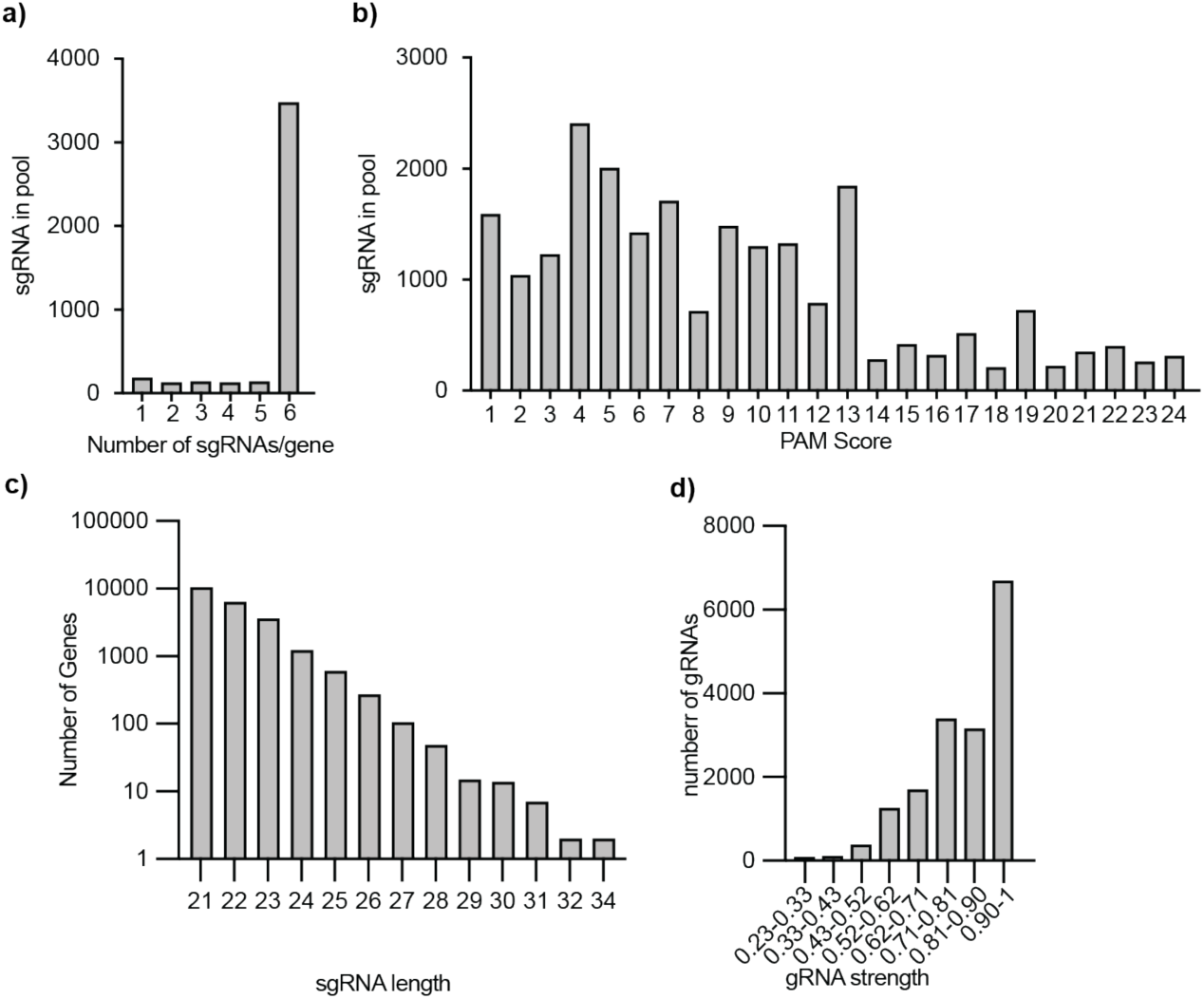
Characteristics of the constructed WG-CRISPRi library. Distribution of **(a)** the number of gRNAs targeting each gene, **(b)** the predicted PAM score of each gRNA based on Rock et al 2017, **(c)** the length of gRNAs and **(d)** the predicted gRNA strength based on Bosch et al 2021.

**Extended Data Fig. 2.**
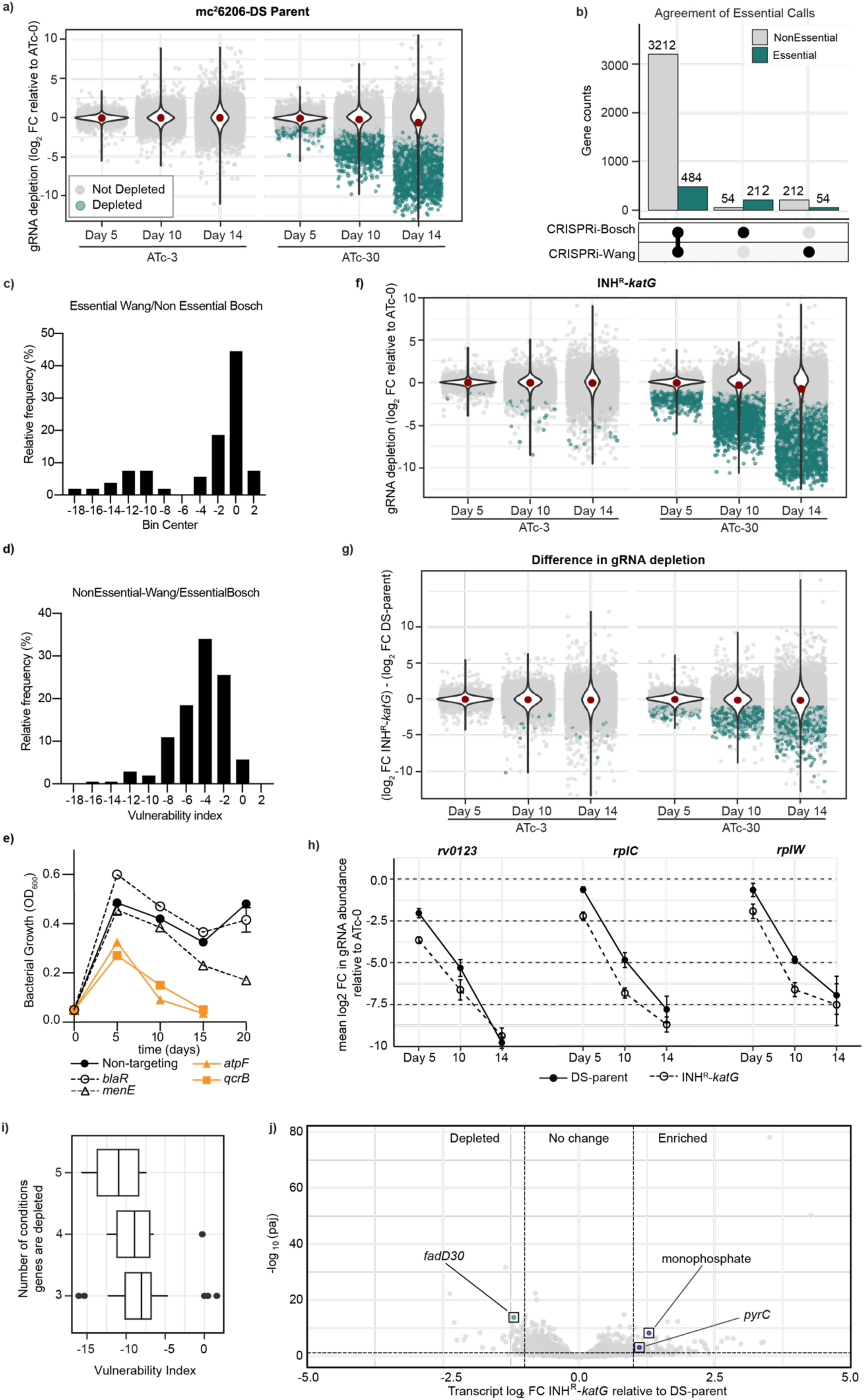
Validation of WG-CRISPRi screening to identify genetic vulnerability in *M. tuberculosis*. **(a)** The summary of gRNA abundance at ATc-3 and ATc-30 in the *M. tuberculosis* strain mc^2^6206 DS-parent. **(b)** The overlap in gene essentiality calls between CRISPRi-Bosch and our WG-CRISPRi study (CRISPRi-Wang). **(c-d)** Distribution of VI of target genes with disagreed essentiality calls between CRISPRi-Bosch and CRISPRi-Wang. Genes frequencies were shown in percentage format (c) VI of disagreed target genes that we called essential but Bosch called non-essential (n = 212) (d)) VI of disagreed target genes that we called non-essential but Bosch called essential (54) **(e)** Growth kinetics of *M. tuberculosis* DS-parent expressing gRNAs that target *menE* and *blaR*, i.e. genes identified as essential in CRISPRi-Bosch, but non-essential in CRISPRi-Wang. A non-targeting gRNA is included as a negative control, whilst gRNA targeting *atpF* and *qcrB* are included as essential genes. Bacterial growth was measured by OD_600_ and back diluted 1/20 into fresh media on days 5, 10, 15 and 20. All strains were grown in 7H9-K with ATc-300. **(f-g)** The summary of gRNA abundance at ATc-3 and ATc-30 in the *M. tuberculosis* strain (f) INH^R^-*katG*, and (g) the depletion difference between the DS-parent and INH^R^-*katG*. **(h)** Mean gRNA abundance targeting genes (Rv0123, *rplC* and *rplW*) that were significantly depleted under no less than 3 conditions but not on day 14 ATc-300. Data were plotted as the changes in gRNA abundance relative to its ATc-0 along time points. **(i)** Summary of genes (n =38) that were more depleted under >= 3 conditions but not on day 14+ATc-300. Genes are grouped by the number of depleted conditions and vulnerability index of each gene from CRISPRi-Bosch is plotted on the x-axis.**(j)** Differentially expressed genes in INH^R^-*katG*. Each dot represents a single gene. Only the differentially expressed and more vulnerable genes in INH^R^-*katG* are coloured and labelled.

**Extended Data Fig. 3.**
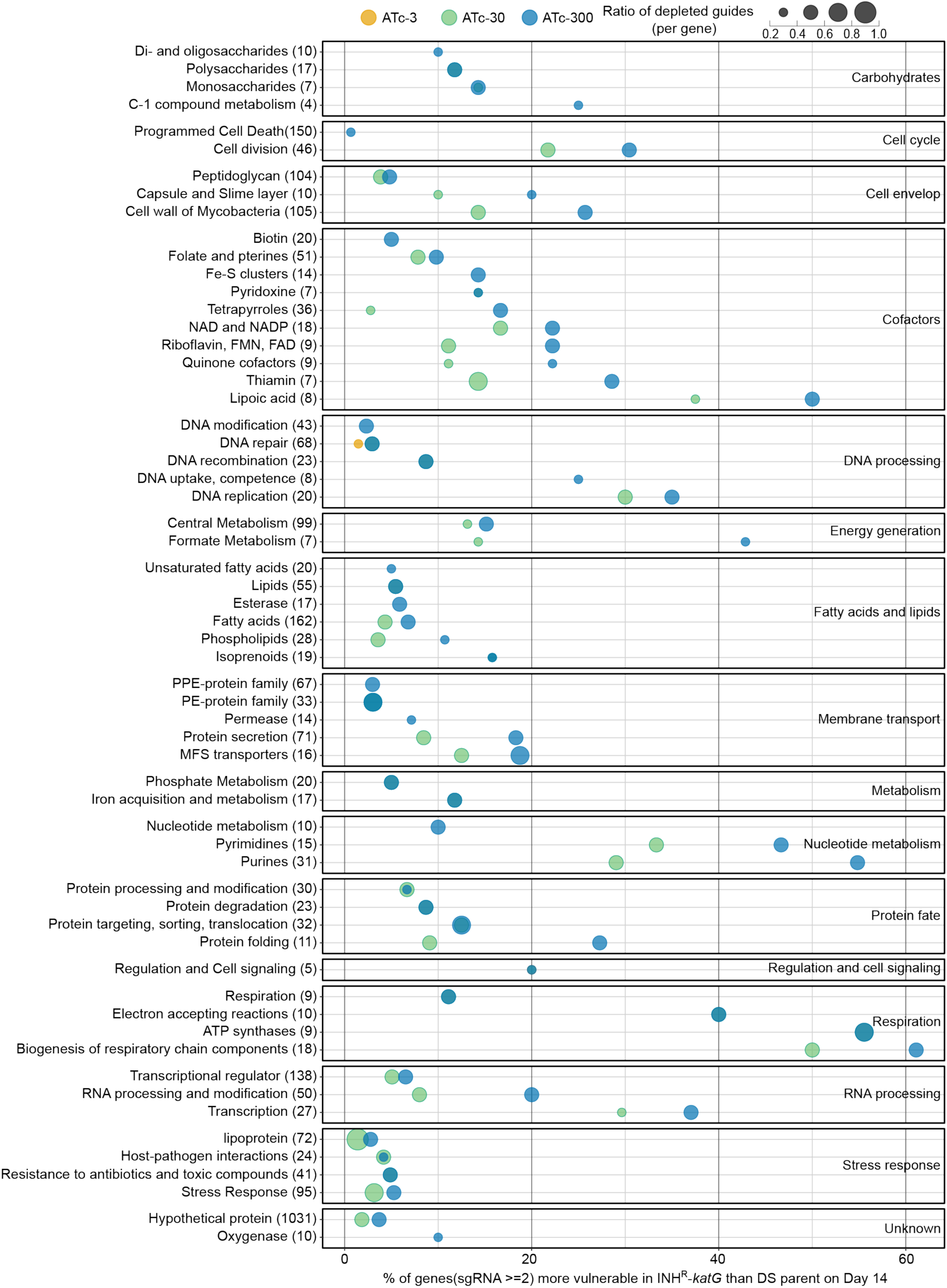
Pathway enrichment analysis of more vulnerable genes in in INH^R^-*katG* as defined by PATRIC functional subclass. Pathway enrichment analysis of genes that are of more vulnerable to inhibition in INH^R^-*katG* according to functional subclass. Each panel represents a PATRIC functional class (shown as the annotated text within each panel) broken down into functional subclass classification. Bubble plot represents data from day 14. Within each functional subclass the size of the bubble indicates the average ratio of gRNAs targeting each gene that is more depleted in INH^R^-*katG.* The colour denotes the ATc concentration from which the amplicon sequencing was performed. The number of genes in each functional class is labelled on the y-axis in brackets. Data for amino acids and derivatives, and protein synthesis functional subclasses is presented in Figures 4 and 5 respectively.

**Extended Data Fig. 4.**
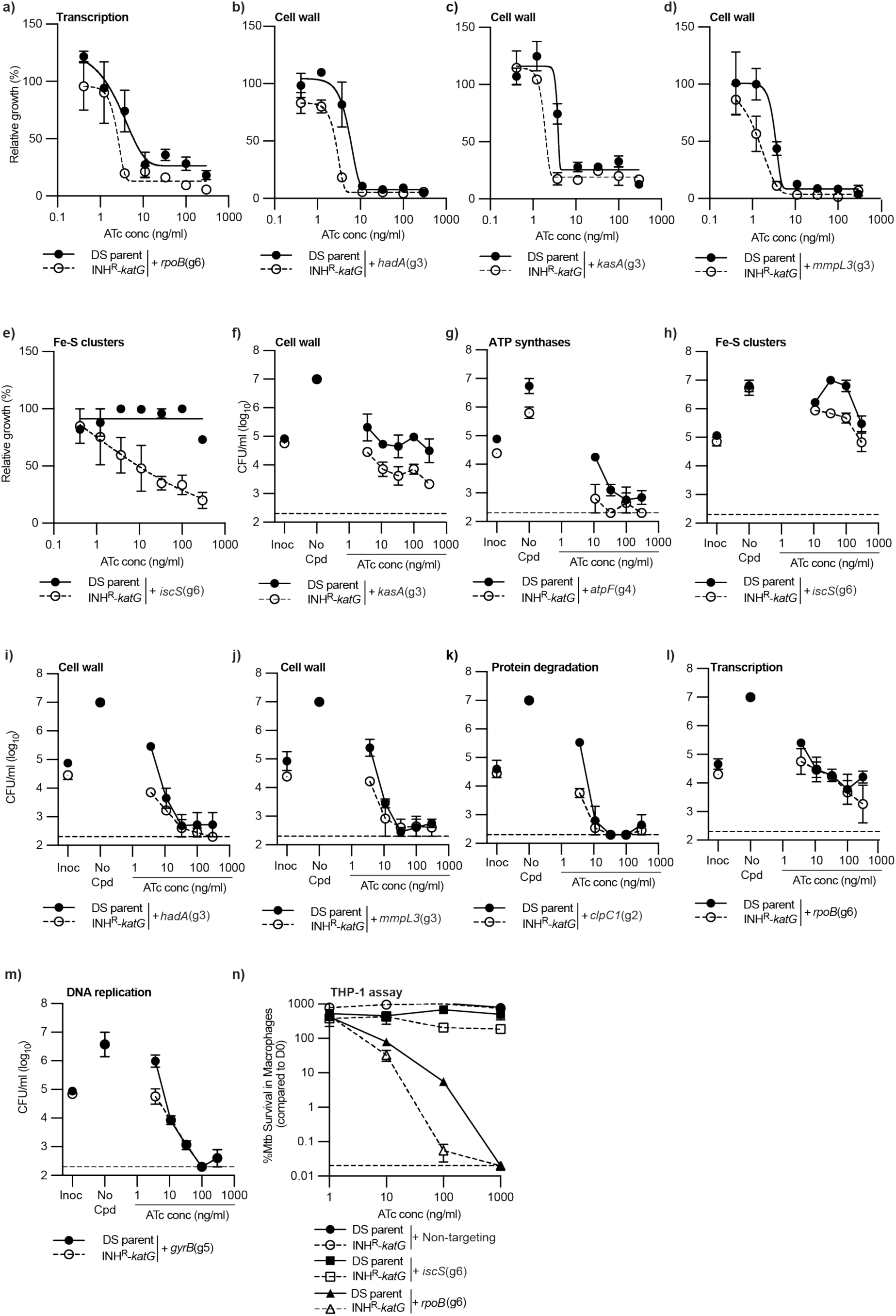
Pathway enrichment analysis defines diverse pathways that are more vulnerable to inhibition in INH^R^-*katG*. **(a-e)** Growth of *M. tuberculosis* DS-parent and INH^R^-*katG* expressing gRNAs that target (a) *rpoB*, (b) *hadA,* (c) *kasA,* (d) *mmpl3* and (e) *iscS* in ATc dose response assays (mean ± range of two biological replicates, n≥3). The (gx) after each gRNA denotes the specific gRNA targeting each gene. **(f-m)** Viability plots of *M. tuberculosis* DS-parent and INH^R^-*katG* expressing for gRNA targeting (f) *kasA,* (g) *atpF,* (h) *iscS,* (i) *hadA,* (j) *mmpl3,* (k) *clpC1*, (l) *rpoB*, and (m) *gyrB.* CFU/ml were determined from 96 well plates at the stated ATc concentrations. Inoc denotes the starting CFU/ml and no-cpd denotes the detected CFU/ml in the absence of ATc (mean ± range of two biological replicates, n≥3). Dashed line represents the lower limit of detection. **(n)** THP-1 macrophage cells were infected with *M. tuberculosis* DS-parent and INH^R^-*katG* cells expressing gRNAs targeting *iscS*, *rpoB* or with a non-targeting control. CRISPRi was induced with the stated concentrations of ATc and intracellular survival was determined by plating for viable colonies (mean ± SD of three biological replicates, n=2).

**Extended Data Fig. 5.**
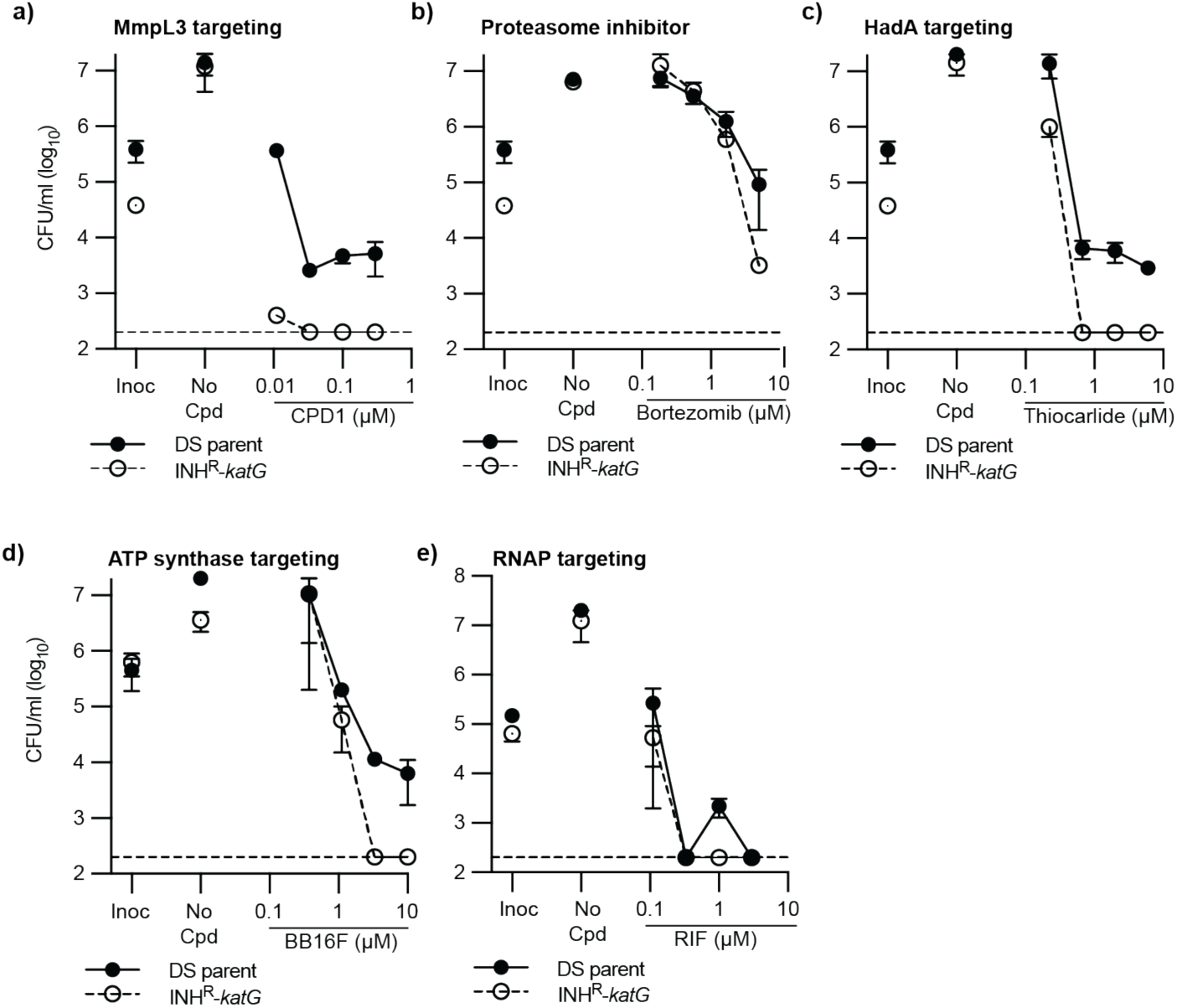
Diverse pathways are more vulnerable to chemical inhibition in INH^R^-*katG*. **(a-e)** MBC assays were used to determine the susceptibility of *M. tuberculosis* DS-parent and INH^R^-*katG* to increasing concentrations of (a) CPD1, (b) Bortezomib, (c) thiocarlide, (d) BB16F and (e) rifampicin. Inoc denotes the starting CFU/ml and no-cpd denotes the detected CFU/ml in the absence of compound (mean ± range of two biological replicates, n≥3). Dashed line represents the lower limit of detection.

**Extended Data Fig. 6.**
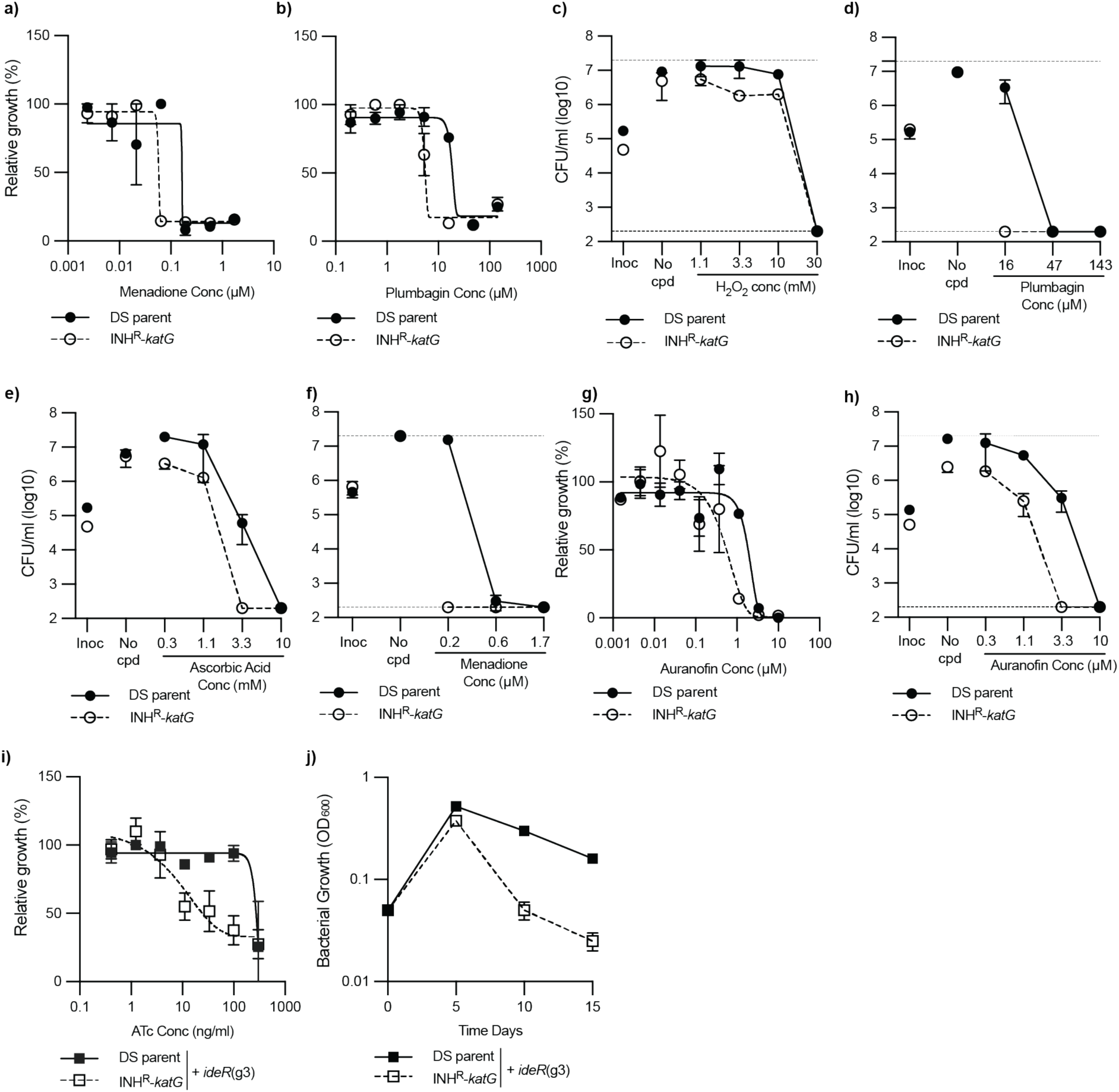
INH^R^-*katG* utilizes alternative redox detoxification pathways to compensate for the loss of *katG*. **(a-b)** Susceptibility of *M. tuberculosis* DS-parent and INH^R^-*katG* to growth inhibition by (a) menadione and (b) plumbagin. **(c-f)**. Susceptibility of *M. tuberculosis* DS-parent and INH^R^-*katG* to killing by (c) H_2_O_2_, (d) plumbagin, (e) ascorbic acid, and (f) menadione. **(g-h)** Susceptibility of *M. tuberculosis* DS-parent and INH^R^-*katG* to auranofin in 96 well plate dose response assays as detected by (g) growth inhibition and (h) killing (mean ± range of two biological replicates, n≥3). Inoc denotes the starting CFU/ml and no-cpd denotes the detected CFU/ml in the absence of compound. Dashed line represents the lower limit of detection. **(i-j)** Growth of *M. tuberculosis* DS-parent and INH^R^-*katG* expressing gRNAs that target *ideR* as detected using (i) 96 well plate ATc dose response assays and (j) by maintaining continuous log-phase growth by back diluting 1/20 into fresh media on days 5, 10, 15. All strains were grown in 7H9-K with ATc-300 and growth was determined by measuring OD_600_.

**Extended Data Fig. 7.**
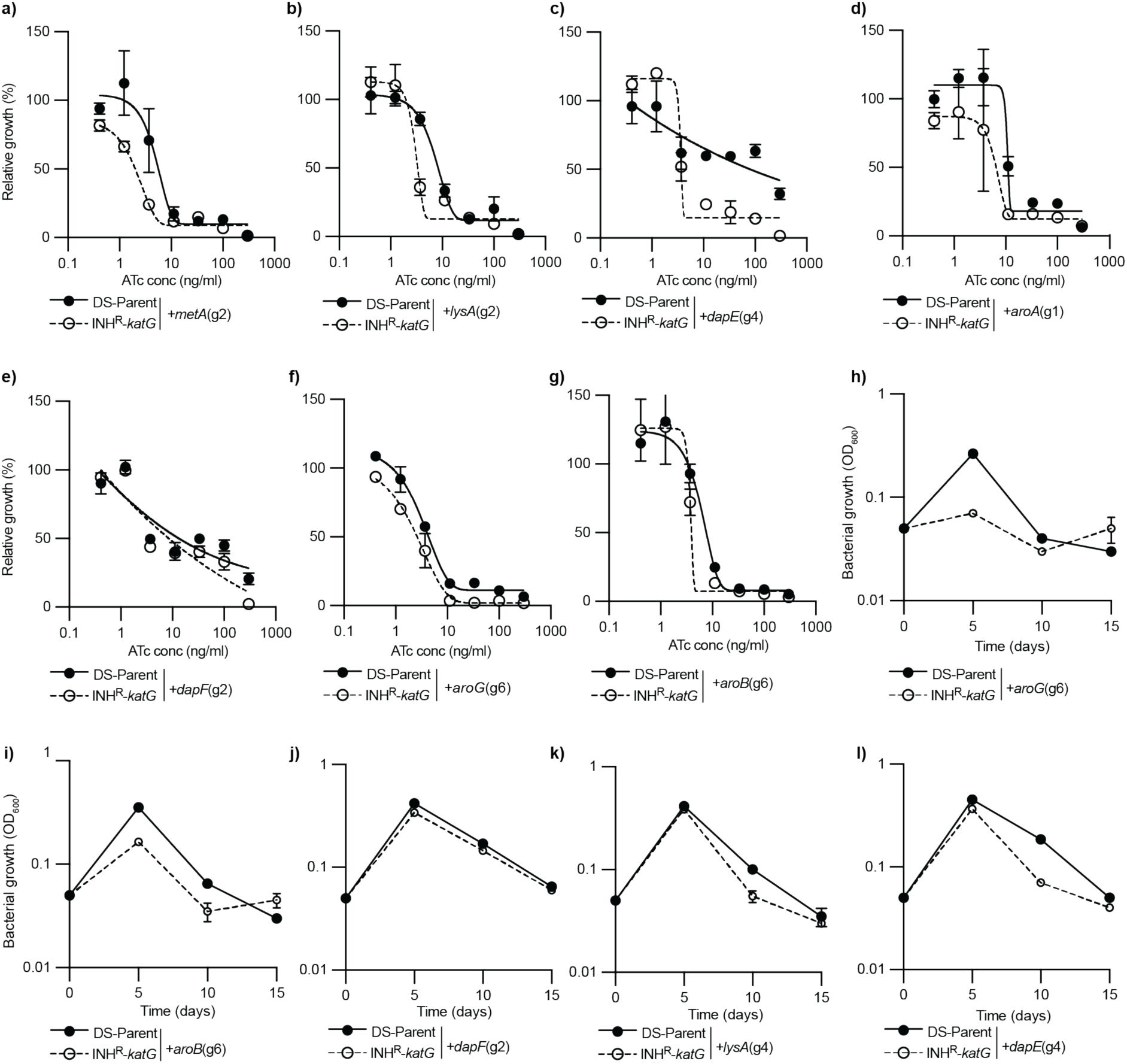
Amino acid metabolism is altered to compensate for the loss of a functional *katG* in *M. tuberculosis*. **(a-g)** Growth of *M. tuberculosis* DS-parent and INH^R^-*katG* expressing gRNA targeting (a) *metA*, (b) *lysA*, (c) *dapE*, (d) *aroA*, (e) *dapF*, (f) *aroG*, and (g) *aroB* in ATc dose response assays (mean ± range of two biological replicates, n≥3). The (gx) after each gRNA denotes the specific gRNA targeting each gene. **(h-l)** Growth kinetics of *M. tuberculosis* DS-parent and INH^R^-*katG* expressing a gRNA targeting (h) *aroG*, (i) aroB, (j) *dapF*, (k) *lysA*, and (l) *dapE*. Bacterial growth was measured by OD_600_ and back diluted 1/20 into fresh media on days 5, 10, 15. All strains were grown in 7H9-K ATc-300.

**Extended Data Fig. 8.**
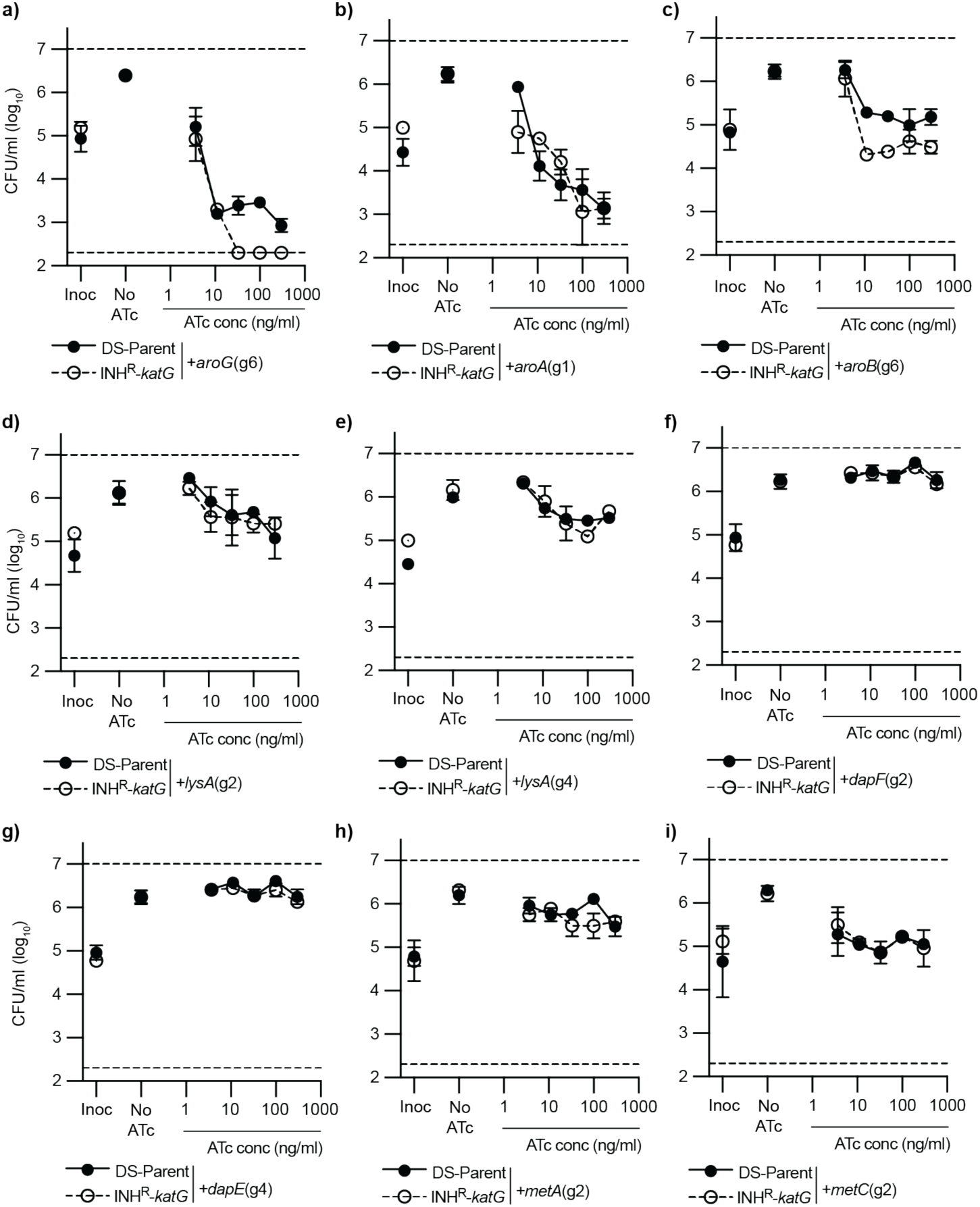
Amino acid metabolism is altered to compensate for the loss of a functional *katG* in *M. tuberculosis*. **(a-i)** Viability plots of *M. tuberculosis* DS-parent and INH^R^-*katG* expressing for gRNA targeting (a) *aroG*, (b) *aroA*, (c) *aroB*, (d-e) *lysA*, (f) *dapF*, (g) *dapE*, (h) *metA*, and (i) *metC*. CFU/ml was determined from 96 well plates. Inoc denotes the starting CFU/ml and no-cpd denotes the detected CFU/ml in the absence of ATc (mean ± range of two biological replicates, n≥3). Dashed line represents the lower limit of detection.

**Extended Data Fig. 9.**
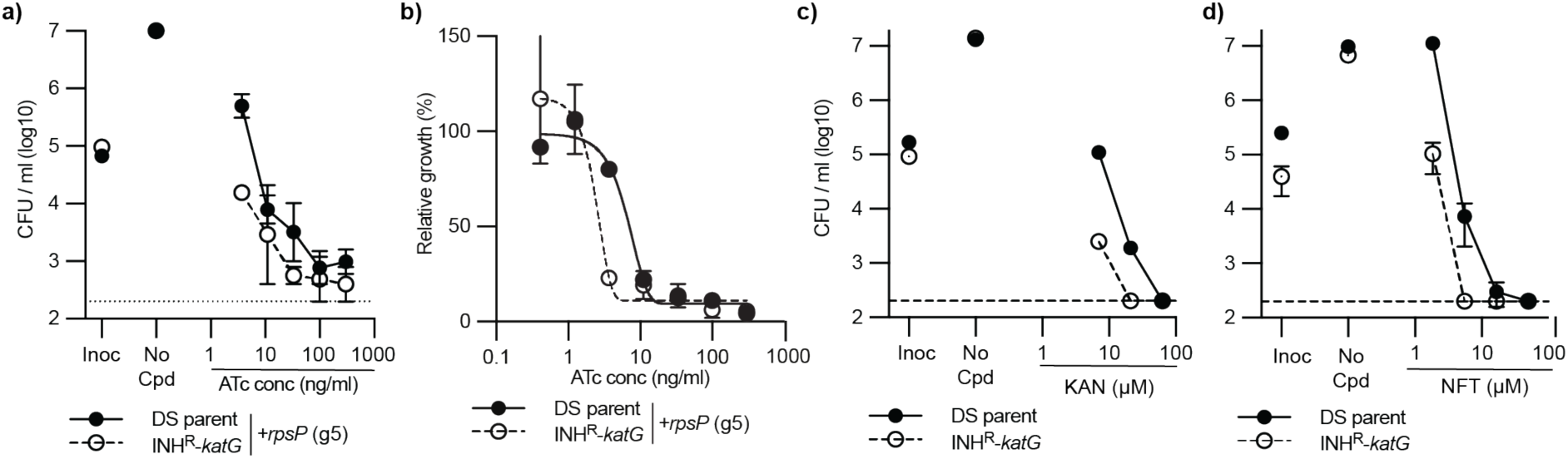
Ribosome biogenesis is more vulnerable to inhibition in an INH^R^-*katG* mutant. **(a)** Viability plots of *M. tuberculosis* DS-parent and INH^R^-*katG* expressing for gRNA targeting *rpsP.* CFU/ml were determined from 96 well plates at the stated ATc concentrations.**(b)** Growth of *M. tuberculosis* DS-parent and INH^R^-*katG* expressing gRNAs targeting *rpsP* in ATc dose response assays (mean ± range of two biological replicates, n≥3). The (gx) after each gRNA denotes the specific gRNA targeting each gene. **(c-d)** MBC assays were used to determine the susceptibility of *M. tuberculosis* DS-parent and INH^R^-*katG* to increasing concentrations of (b) Kanamycin and (c) nitrofurantoin. For a-c Inoc denotes the starting CFU/ml and no-cpd denotes the detected CFU/ml in the absence of compound or ATc (mean ± range of two biological replicates, n≥3). Dashed line represents the lower limit of detection.

## References

1. World Health Organization. WHO TB Report-2022.

2. Conradie F, Diacon AH, Ngubane N, Howell P, Everitt D, Crook AM, et al. Treatment of Highly Drug-Resistant Pulmonary Tuberculosis. New England Journal of Medicine. 2020 Mar 5;382(10):893–902.

3. Pym AS, Saint-Joanis B, Cole ST. Effect of katG mutations on the virulence of Mycobacterium tuberculosis and the implication for transmission in humans. Infect Immun. 2002;70(9):4955–60.

4. Sherman DR, Mdluli K, Hickey MJ, Arain TM, Morris L, Iii CEB, et al. Compensatory ahpC Gene Expression in Isoniazid-Resistant Mycobacterium tuberculosis. Science (1979). 1996;272(5268):1641–3.

5. Zhang Y, Heym B, Allen B, Young D, Cole ST. The catalase-peroxidase gene and isoniazid resistance of *Mycobacterium tuberculosis*. Nature. 1992;358:591–3.

6. Rittershaus ESC, Baek SH, Krieger I V., Nelson SJ, Cheng YS, Nambi S, et al. A Lysine Acetyltransferase Contributes to the Metabolic Adaptation to Hypoxia in Mycobacterium tuberculosis. Cell Chem Biol. 2018 Dec 20;25(12):1495–1505.e3.

7. Zhang YJ, Reddy MC, Ioerger TR, Rothchild AC, Dartois V, Schuster BM, et al. Tryptophan biosynthesis protects mycobacteria from CD4 T-Cell-mediated Killing. Cell. 2013;155(6):1296–308.

8. Parikh SL, Xiao G, Tonge PJ. Inhibition of InhA, the enoyl reductase from Mycobacterium tuberculosis, by triclosan and isoniazid. Biochemistry. 2000;39(26):7645–50.

9. Xia Y, Zhou Y, Carter DS, Mcneil MB, Choi W, Halladay J, et al. Discovery of a cofactor-independent inhibitor of Mycobacterium tuberculosis InhA. 2018;1(3):1–12.

10. Crook DW, Peto TEA, Hoosdally SJ, Cruz ALG, Walker AS, Walker TM, et al. A data compendium associating the genomes of 12,289 Mycobacterium tuberculosis isolates with quantitative resistance phenotypes to 13 antibiotics. PLoS Biol. 2022 Aug 1;20(8):e3001721.

11. Lou K, Steri V, Ge AY, Hwang YC, Yogodzinski CH, Shkedi AR, et al. KRASG12C inhibition produces a driver-limited state revealing collateral dependencies. Sci Signal. 2019;12(583).

12. Wang C, Vegna S, Jin H, Benedict B, Lieftink C, Ramirez C, et al. Inducing and exploiting vulnerabilities for the treatment of liver cancer. Nature. 2019;574:268–72.

13. Wang L, Bernards R. Taking advantage of drug resistance, a new approach in the war on cancer. Front Med. 2018;12(4):490–5.

14. Brandis G, Wrande M, Liljas L, Hughes D. Fitness-compensatory mutations in rifampicin-resistant RNA polymerase. Mol Microbiol. 2012;85(1):142–51.

15. Gagneux S, Long CD, Small PM, Van T, Schoolnik GK, Bohannan BJM. The Competitive Cost of Antibiotic Resistance in *Mycobacterium tuberculosis*. Science (1979). 2015;312(5782):1944–6.

16. Gonzales PR, Pesesky MW, Bouley R, Ballard A, Biddy BA, Suckow MA, et al. Synergistic, collaterally sensitive β-lactam combinations suppress resistance in MRSA. Nat Chem Biol. 2015;11(11):855–64.

17. Barbosa C, Trebosc V, Kemmer C, Rosenstiel P, Beardmore R, Schulenburg H, et al. Alternative evolutionary paths to bacterial antibiotic resistance cause distinct collateral effects. Mol Biol Evol. 2017;1–16.

18. Munck C, Gumpert HK, Wallin AI, Wang HH, Sommer MO. Prediction of resistance development against drug combinations by collateral responses to component drugs. Sci Transl Med. 2014;6(262):262ra156.

19. Waller NJE, Cheung CY, Cook GM, McNeil MB. The evolution of antibiotic resistance is associated with collateral drug phenotypes in Mycobacterium tuberculosis. Nat Commun. 2023 Mar 18;14(1):1517.

20. Rodriguez De Evgrafov M, Gumpert H, Munck C, Thomsen TT, Sommer MOA. Collateral resistance and sensitivity modulate evolution of high-level resistance to drug combination treatment in staphylococcus aureus. Mol Biol Evol. 2015;32(5):1175–85.

21. Imamovic L, Sommer MOA. Use of Collateral Sensitivity Networks to Design Drug Cycling Protocols That Avoid Resistance Development. Sci Transl Med. 2013;5(204).

22. Imamovic L, Mostafa M, Ellabaan H, Manuel A, Machado D, Molin S, et al. Drug-Driven Phenotypic Convergence Supports Rational Treatment Strategies of Chronic Infections. Cell. 2018;172:121–34.

23. Rock JM, Hopkins FF, Chavez A, Diallo M, Chase MR, Gerrick ER, et al. Programmable transcriptional repression in mycobacteria using an orthogonal CRISPR interference platform. Nat Microbiol. 2017;2(February):1–9.

24. Rock J. Tuberculosis drug discovery in the CRISPR era. 2019;1–10.

25. Bosch B, Dejesus MA, Poulton NC, Zhang W, Engelhart CA, Zaveri A, et al. Genome-wide gene expression tuning reveals diverse vulnerabilities of M. tuberculosis. Cell. 2021;184(17):4579–92.

26. Jost M, Santos DA, Saunders RA, Horlbeck MA, Hawkins JS, Scaria SM, et al. Titrating gene expression using libraries of systematically attenuated CRISPR guide RNAs. Nat Biotechnol. 2020 Mar 1;38(3):355–64.

27. JS H. Mismatch-CRISPRi Reveals the Co-varying Expression-Fitness Relationships of Essential Genes in Escherichia coli and Bacillus subtilis. 2020.

28. Donati S, Kuntz M, Pahl V, Randau L, Wang C ying, Link H, et al. Multi-omics Analysis of CRISPRi-Knockdowns Identifies Mechanisms that Buffer Decreases of Enzymes in E. coli Metabolism. Cell Syst. 2021;12(1):56–67.

29. McNeil MB, Keighley LM, Cook JR, Cheung CY, Cook GM. CRISPR interference identifies vulnerable cellular pathways with bactericidal phenotypes in Mycobacterium tuberculosis. Mol Microbiol. 2021 Oct 1;116(4):1033–43.

30. Mcneil MB, Ryburn HWK, Harold LK, Tirados JF, Cook GM. Transcriptional Inhibition of the F 1 F 0-Type ATP Synthase Has Bactericidal Consequences on the Viability of Mycobacteria. Antimicrob Agents Chemother. 2020;64(8).

31. Cheung CY, McNeil MB, Cook GM. Utilization of CRISPR interference to investigate the contribution of genes to pathogenesis in a macrophage model of Mycobacterium tuberculosis infection. Journal of Antimicrobial Chemotherapy. 2022 Mar 1;77(3):615– 9.

32. Sies H, Belousov V V., Chandel NS, Davies MJ, Jones DP, Mann GE, et al. Defining roles of specific reactive oxygen species (ROS) in cell biology and physiology. Nat Rev Mol Cell Biol. 2022 Jul 1;23(7):499–515.

33. Murphy MP, Bayir H, Belousov V, Chang CJ, Davies KJA, Davies MJ, et al. Guidelines for measuring reactive oxygen species and oxidative damage in cells and in vivo. Nat Metab. 2022 Jun 1;4(6):651–62.

34. Vilchèze C, Hartman T, Weinrick B, Jacobs WR. Mycobacterium tuberculosis is extraordinarily sensitive to killing by a vitamin C-induced Fenton reaction. Nat Commun. 2013;4.

35. Vilchèze C, Hartman T, Weinrick B, Jain P, Weisbrod TR, Leung LW, et al. Enhanced respiration prevents drug tolerance and drug resistance in Mycobacterium tuberculosis. Proceedings of the National Academy of Sciences. 2017;114(17):4495– 500.

36. Lin K, Brien KMO, Trujillo C, Wang R, Wallach JB. Mycobacterium tuberculosis Thioredoxin Reductase Is Essential for Thiol Redox Homeostasis but Plays a Minor Role in Antioxidant Defense. 2016;1–20.

37. Reddy PV, Puri RV, Khera A, Tyagi AK. Iron Storage Proteins Are Essential for the Survival and Pathogenesis of Mycobacterium tuberculosis in THP-1 Macrophages and the Guinea Pig Model of Infection. 2012;567–75.

38. Dragset MS, Ioerger TR, Zhang YJ, Mærk M, Ginbot Z, Sacchettini JC, et al. Genome-wide Phenotypic Profiling Identifies and Categorizes Genes Required for Mycobacterial Low Iron Fitness. Sci Rep. 2019 Dec 1;9(1).

39. Theriault ME, Pisu D, Wilburn KM, Lê-Bury G, MacNamara CW, Michael Petrassi H, et al. Iron limitation in M. tuberculosis has broad impact on central carbon metabolism. Commun Biol. 2022 Dec 1;5(1).

40. Rodriguez GM, Voskuil MI, Gold B, Schoolnik GK, Smith I. ideR, an Essential Gene in Mycobacterium tuberculosis : Role of IdeR in Iron-Dependent Gene Expression, Iron Metabolism, and Oxidative Stress Response. Infect Immun. 2002;70(7):3371– 81.

41. Pandey R, Rodriguez GM. IdeR is required for iron homeostasis and virulence in Mycobacterium tuberculosis. Mol Microbiol. 2014 Jan;91(1):98–109.

42. Ofori-Anyinam N, Hamblin M, Coldren M, Li B, Mereddy G, Shaikh M, et al. KatG catalase deficiency confers bedaquiline hyper-susceptibility to isoniazid resistant Mycobacterium tuberculosis. bioRxiv. 2023 Oct 17;

43. Trauner A, Lougheed KEA, Bennett MH, Hingley-Wilson SM, Williams HD. The dormancy regulator DosR controls ribosome stability in hypoxic mycobacteria. Journal of Biological Chemistry. 2012;287(28):24053–63.

44. Mehra S, Foreman TW, Didier PJ, Ahsan MH, Hudock T a, Kissee R, et al. The DosR Regulon Modulates Adaptive Immunity and is Essential for M. tuberculosis Persistence. Am J Respir Crit Care Med. 2015;191(10):1185–96.

45. He H, Bretl DJ, Penoske RM, Anderson DM, Zahrt TC. Components of the Rv0081-Rv0088 Locus, which encodes a predicted formate hydrogenlyase complex, are coregulated by Rv0081, MprA, and DosR in Mycobacterium tuberculosis. J Bacteriol. 2011 Oct;193(19):5105–18.

46. Roberts DM, Liao RP, Wisedchaisri G, Hol WGJ, Sherman DR. Two sensor kinases contribute to the hypoxic response of Mycobacterium tuberculosis. Journal of Biological Chemistry. 2004;279(22):23082–7.

47. Kumar A, Toledo JC, Patel RP, Lancaster JR, Steyn AJC, Designed AJCS, et al. Mycobacterium tuberculosis DosS is a redox sensor and DosT is a hypoxia sensor. Vol. 104. 2007.

48. Zheng H, Colvin CJ, Johnson BK, Kirchhoff PD, Wilson M, Jorgensen-Muga K, et al. Inhibitors of Mycobacterium tuberculosis DosRST signaling and persistence. Nat Chem Biol. 2017 Feb 1;13(2):218–25.

49. Kung-Chun Chiu D, Pui-Wah Tse A, Law CT, Ming-Jing Xu I, Lee D, Chen M, et al. Hypoxia regulates the mitochondrial activity of hepatocellular carcinoma cells through HIF/HEY1/PINK1 pathway. Cell Death Dis. 2019 Dec 1;10(12).

50. Coimbra-Costa D, Alva N, Duran M, Carbonell T, Rama R. Oxidative stress and apoptosis after acute respiratory hypoxia and reoxygenation in rat brain. Redox Biol. 2017 Aug 1;12:216–25.

51. Turrens JF. Mitochondrial formation of reactive oxygen species. Vol. 552, Journal of Physiology. 2003. p. 335–44.

52. Chandel NS, Maltepe E, Goldwasser E, Mathieu CE, Simon MC, Schumacker PT. Mitochondrial reactive oxygen species trigger hypoxia-induced transcription. PNAS. 1998;95(20):11715–20.

53. Mackenzie JS, Lamprecht DA, Asmal R, Adamson JH, Borah K, Beste DJ V, et al. Bedaquiline reprograms central metabolism to reveal glycolytic vulnerability in Mycobacterium tuberculosis. 2020;

54. Eoh H, Rhee KY. Multifunctional essentiality of succinate metabolism in adaptation to hypoxia in Mycobacterium tuberculosis. Proc Natl Acad Sci U S A. 2013 Apr 16;110(16):6554–9.

55. Ling J, Söll D. Severe oxidative stress induces protein mistranslation through impairment of an aminoacyl-tRNA synthetase editing site. Proc Natl Acad Sci U S A. 2010 Mar 2;107(9):4028–33.

56. Kohanski MA, Dwyer DJ, Hayete B, Lawrence CA, Collins JJ. A Common Mechanism of Cellular Death Induced by Bactericidal Antibiotics. Cell. 2007 Sep 7;130(5):797– 810.

57. Dwyer DJ, Kohanski MA, Collins JJ. Role of reactive oxygen species in antibiotic action and resistance. Curr Opin Microbiol. 2009;12(5):482–9.

58. Dwyer DJ, Collins JJ, Walker GC. Unraveling the Physiological Complexities of Antibiotic Lethality. Annu Rev Pharmacol Toxicol. 2015;55(1):313–32.

59. Belenky P, Ye JD, Porter CBM, Cohen NR, Lobritz MA, Ferrante T, et al. Bactericidal Antibiotics Induce Toxic Metabolic Perturbations that Lead to Cellular Damage. Cell Rep. 2015;13(5):968–80.

60. Hong Y, Zeng J, Wang X, Drlica K, Zhao X. Post-stress bacterial cell death mediated by reactive oxygen species. Proc Natl Acad Sci U S A. 2019;116(20):10064–71.

61. Wang X, Zhao X. Contribution of oxidative damage to antimicrobial lethality. Antimicrob Agents Chemother. 2009 Apr;53(4):1395–402.

62. Olin-Sandoval V, Yu JSL, Miller-Fleming L, Alam MT, Kamrad S, Correia-Melo C, et al. Lysine harvesting is an antioxidant strategy and triggers underground polyamine metabolism. Nature. 2019 Aug 8;572(7768):249–53.

63. Loots DT. An altered Mycobacterium tuberculosis metabolome induced by katG mutations resulting in isoniazid resistance. Antimicrob Agents Chemother. 2014;58(4):2144–9.

64. Hasenoehrl EJ, Rae Sajorda D, Berney-Meyer L, Johnson S, Tufariello JAM, Fuhrer T, et al. Derailing the aspartate pathway of Mycobacterium tuberculosis to eradicate persistent infection. Nat Commun. 2019 Dec 1;10(1).

65. Berney M, Berney-Meyer L. Mycobacterium tuberculosis in the Face of Host-Imposed Nutrient Limitation. Microbiol Spectr. 2017;5(3):1–17.

66. Berney M, Berney-Meyer L, Wong KW, Chen B, Chen M, Kim J, et al. Essential roles of methionine and S-adenosylmethionine in the autarkic lifestyle of Mycobacterium tuberculosis. Proceedings of the National Academy of Sciences. 2015;112(32):10008–13.

67. Tiwari S, Van Tonder AJ, Vilchèze C, Mendes V, Thomas SE, Malek A, et al. Arginine-deprivation-induced oxidative damage sterilizes Mycobacterium tuberculosis. Proc Natl Acad Sci U S A. 2018;115(39):9779–84.

68. Kumar A, Farhana A, Guidry L, Saini V, Hondalus M, Steyn AJC. Redox homeostasis in mycobacteria: the key to tuberculosis control? Expert Rev Mol Med. 2011;13(December):1–25.

69. Singh A, Crossman DK, Mai D, Guidry L, Voskuil MI, Renfrow MB, et al. Mycobacterium tuberculosis WhiB3 Maintains redox homeostasis by regulating virulence lipid anabolism to modulate macrophage response. PLoS Pathog. 2009 Aug;5(8).

70. Saini V, Cumming BM, Guidry L, Lamprecht DA, Adamson JH, Reddy VP, et al. Ergothioneine Maintains Redox and Bioenergetic Homeostasis Essential for Drug Susceptibility and Virulence of Mycobacterium tuberculosis. Cell Rep. 2016;14(3):572–85.

71. Peters JM, Colavin A, Shi H, Czarny TL, Larson MH, Wong S, et al. A Comprehensive, CRISPR-based Functional Analysis of Essential Genes in Bacteria. Cell. 2016;165(6):1493–506.

72. Qu J, Prasad NK, Yu MA, Chen S, Lyden A, Herrera N, et al. Modulating Pathogenesis with Mobile-CRISPRi. J Bacteriol. 2019;201(22):00304–19.

73. Peters JM, Koo BM, Patino R, Heussler GE, Hearne CC, Qu J, et al. Enabling genetic analysis of diverse bacteria with Mobile-CRISPRi. Vol. 4, Nature Microbiology. Nature Publishing Group; 2019. p. 244–50.

74. Jain P, Hsu T, Arai M, Biermann K, Thaler DS, Nguyen A, et al. Specialized Transduction Designed for Precise High-Throughput Unmarked Deletions in Mycobacterium tuberculosis. 2014;5(3):1–9.

75. DeJesus MA, Gerrick ER, Xu W, Park SW, Long JE, Boutte CC, et al. Comprehensive Essentiality Analysis of the Mycobacterium tuberculosis Genome via. mBio. 2017;8(1):1–17.

76. McNeil MB, Cook GM. Utilization of CRISPR interference to validate MmpL3 as a drug target in mycobacterium tuberculosis. Antimicrob Agents Chemother. 2019;63(8).

77. Shen W, Le S, Li Y, Hu F. SeqKit: A cross-platform and ultrafast toolkit for FASTA/Q file manipulation. PLoS One. 2016 Oct 1;11(10).

78. Chen Y, Lun ATL, Smyth GK. From reads to genes to pathways: differential expression analysis of RNA-Seq experiments using Rsubread and the edgeR quasi-likelihood pipeline. F1000Res. 2016 Jun 20;5:1438.

79. Gillespie JJ, Wattam AR, Cammer SA, Gabbard JL, Shukla MP, Dalay O, et al. Patric: The comprehensive bacterial bioinformatics resource with a focus on human pathogenic species. Infect Immun. 2011 Nov;79(11):4286–98.

80. McNeil MB, Ryburn HW, Tirados J, Cheung CY, Cook GM. Multiplexed transcriptional repression identifies a network of bactericidal interactions between mycobacterial respiratory complexes. iScience. 2022 Jan 21;25(1).

81. Hards K, Cheung CY, Waller N, Adolph C, Keighley L, Tee ZS, et al. An amiloride derivative is active against the F1Fo-ATP synthase and cytochrome bd oxidase of Mycobacterium tuberculosis. Commun Biol. 2022 Dec 1;5(1).

82. McNeil MB, Dennison DD, Shelton CD, Parish T. In vitro isolation and characterization of oxazolidinone-resistant Mycobacterium tuberculosis. Antimicrob Agents Chemother. 2017;61(10).

83. Kalia NP, Singh S, Hards K, Cheung CY, Sviriaeva E, Banaei-Esfahani A, et al. M. tuberculosis relies on trace oxygen to maintain energy homeostasis and survive in hypoxic environments. Cell Rep. 2023 May 30;42(5).

84. Fridianto KT, Li M, Hards K, Negatu DA, Cook GM, Dick T, et al. Functionalized Dioxonaphthoimidazoliums: A Redox Cycling Chemotype with Potent Bactericidal Activities against Mycobacterium tuberculosis. J Med Chem. 2021 Nov 11;64(21):15991–6007.

85. Masukagami Y, Nijagal B, Mahdizadeh S, Tseng CW, Dayalan S, Tivendale KA, et al. A combined metabolomic and bioinformatic approach to investigate the function of transport proteins of the important pathogen Mycoplasma bovis. Vet Microbiol. 2019 Jul 1;234:8–16.

86. Pang Z, Chong J, Zhou G, De Lima Morais DA, Chang L, Barrette M, et al. MetaboAnalyst 5.0: Narrowing the gap between raw spectra and functional insights. Nucleic Acids Res. 2021 Jul 2;49(W1):W388–96.

87. Bushnell B. BBMap: A Fast, Accurate, Splice-Aware Aligner. Lawrence Berkeley National Laboratory. In 2014.

88. Li H, Handsaker B, Wysoker A, Fennell T, Ruan J, Homer N, et al. The Sequence Alignment/Map format and SAMtools. Bioinformatics. 2009 Aug;25(16):2078–9.

89. Morgan M, Pagès H, Obenchain V, Hayden N. Rsamtools: Binary alignment (BAM), FASTA, variant call (BCF), and tabix file import. R package version 2.16.0. 2023.

90. Lawrence M, Huber W, Pagès H, Aboyoun P, Carlson M, Gentleman R, et al. Software for Computing and Annotating Genomic Ranges. PLoS Comput Biol. 2013;9(8).

91. The CRyPTIC Consortium. A data compendium associating the genomes of 12,289 Mycobacterium tuberculosis isolates with quantitative resistance phenotypes to 13 antibiotics. PLoS Biol. 2022 Aug 1;20(8):e3001721.

